# Quantifying phage infectivity from characteristics of bacterial population dynamics

**DOI:** 10.1101/2023.06.29.546975

**Authors:** Michael Blazanin, Eli Vasen, Cèlia Vilaró Jolis, William An, Paul E. Turner

## Abstract

A frequent goal of phage biology is to quantify how well a phage kills a population of host bacteria. Unfortunately, traditional methods to quantify phage success can be time-consuming, limiting the throughput of experiments. Here, we use theory to show how the effects of phages on their hosts can be quantified using bacterial population dynamics measured in a high-throughput microplate reader (automated spectrophotometer). We use mathematical models to simulate bacterial population dynamics where specific phage and bacterial traits are known *a priori*. We then test common metrics of those dynamics (e.g. growth rate, time and height of peak bacterial density, death rate, extinction time, area under the curve) to determine which best predict: 1) infectivity over the short-term, and 2) phage suppression over the long-term. We find that many metrics predict infectivity and are strongly correlated with one another. We also find that metrics can predict phage growth rate, providing an effective way to quantify the combined effects of multiple phage traits. Finally, we show that peak density, time of peak density, and extinction time are the best metrics when comparing across different bacterial hosts or over longer timescales where plasticity or evolution may play a role. In all, we establish a foundation for using bacterial population dynamics to quantify the effects of phages on their bacterial hosts, supporting the design of *in vitro* empirical experiments using microplate readers.

**Significance:** Bacteriophages are viruses that infect bacteria, with relevance from basic science to medical application. Frequently we seek to quantify how these viruses negatively impact bacterial growth. Typical methods are labor-intense, limiting the number of experiments that can be done. Here, we show how easily-collectable data (called ‘bacterial population dynamics’ or ‘growth curves’) can be used to quantify virus killing of bacteria across a wide range of conditions. In all, our work suggests that these dynamics provide an effective and high-throughput method to quantify phage effects on their hosts.

## Introduction

In the study of lytic bacteriophages (phages), a common experimental goal is to quantify how well a phage can kill a given bacterial strain. This question is asked both on short timescales as well as over longer timescales. On short timescales we want to quantify ‘infectivity’, which we define as how well a specific phage strain infects and kills a specific bacterial strain. At this timescale, we expect that phage and bacterial phenotypes are mainly constant, and that infectivity reflects both: 1) the phage’s ability to infect host cells, as well as 2) the bacterium’s ability to resist phage attack (1, 2). On longer timescales, plasticity and evolution can come into play, allowing phage and bacterial phenotypes to change. In this case, we want to quantify the magnitude and duration of phage-driven suppression of the bacterial population, even as bacteria become resistant to phages. Unfortunately, experimentally quantifying phage success over the short or long-term can require time-consuming methods, limiting the throughput of experiments (3–8).

Here, we aim to show how the success of obligately-lytic phages over both the short and long-term can be quantified in high-throughput from bacterial population dynamics estimated in a microplate reader (‘growth curves’, e.g. as measured with optical density). Many groups have measured proxies of bacterial density over time in the presence and absence of phages in order to qualitatively infer phage activity (9–36). In addition, some studies have proposed ways to use population dynamics to quantify infectivity (37–50), attempting to extract a metric (or metrics) from the time series of bacterial densities that reflects infectivity. However, which of these proposed approaches best reflect infectivity, and whether they can be extended to quantify phage success over longer timescales with plasticity or evolution, has not been systematically tested.

To address this gap, we used mathematical models to simulate bacterial population dynamics in the presence and absence of infecting phages (Fig S1). Such models have been widely used to study phage-bacteria interactions (51–53), but only rarely applied to understand how population dynamics could be used to quantify phage success (45, 47). By modeling, we can simulate and compare bacterial population dynamics where bacterial and phage traits are precisely controlled and known *a priori*. Using our models, we complement the existing body of largely-empirical papers (37–50) to show that many metrics predict infectivity, are strongly correlated with one another, and can be used to infer phage growth rate, providing an effective way to quantify the combined effects of multiple phage traits. Additionally, we find that peak density, time of peak density, and extinction time are the best metrics when bacterial hosts vary in their growth or over longer timescales where plasticity or evolution may play a role.

## Materials and Methods

### Overview

We used the common approach of modeling populations of susceptible bacteria, infected bacteria, free phage particles, and nutrients with a system of differential equations (Fig S1). These equations track the densities of each of the four populations over time, linking the effects of each population on the others through mathematical terms for processes like growth, infection, and lysis. Such terms also include parameters like growth rate, stationary phase density, infection rate, burst size (number of phages produced by each infected cell), and lysis time (lag between infection of a susceptible cell and lysis to release new phage particles). By simulating population dynamics with different parameter values, we can then observe how changes in each parameter value alter the observed bacterial population dynamics. Finally, we can apply the various approaches to analyze growth curves from the literature to our simulated data, enabling us to directly test the efficacy of each approach.

### Main model of bacterial population dynamics

We built a delay differential equation model of bacterial and phage growth (54) (Eqs. 1 – 4) with populations of susceptible bacteria (*S*), infected bacteria (*I*), free phages (*P*), and nutrients (*N*). The nutrient population is defined in units of cells, so that one unit of nutrients yields one cell. In this model, nutrients can return into the environment from cell lysis at a rate controlled by the *d* parameter. Note that when *d* = 1 (i.e. the lysis of one cell returns one cell’s-worth of nutrients back into the environment), this model simplifies to logistic growth. For convenience, we let k be the stationary phase density of cells, and let *N*_0_ = *k* − *S*_0_ − *I*_0_. Subscripts denote time (i.e. *S*_*t*_ is the population of susceptible cells at time t). Descriptions of all parameters are provided in Table 1.

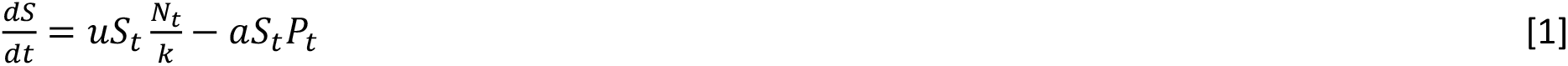

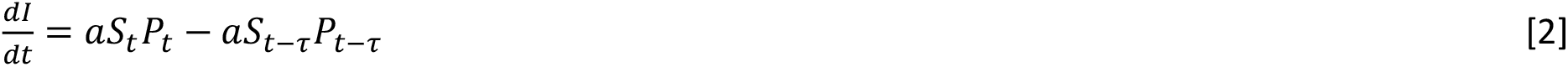

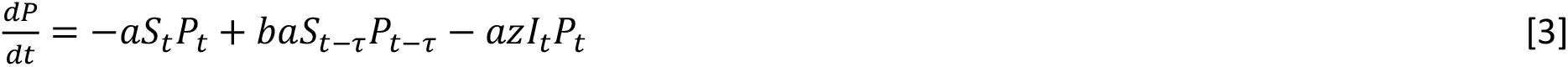

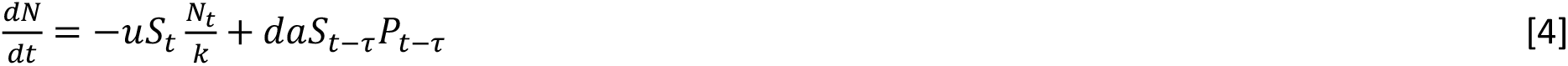

**Table 1.** Model parameter terms and their definitions. Unless otherwise noted, all simulations used the default value listed. For the noted exceptions, simulations typically used the range of values in brackets, which approximately span the experimentally-observed ranges of phenotypes (Table S1).

| Parameter | Definition | Default Value [Range of values] |
| --- | --- | --- |
| $u_S$ | Maximum per-capita growth rate of susceptible cells | 0.0179 [0.00798 – 0.04] /min<br>(doubling time of ~39 [87 – 17] minutes) |
| $k$ | Stationary phase density of cells | $10^9$ [$10^8$ – $10^{10}$] CFU/mL |
| $a$ | Maximum infection rate of susceptible cells by free phage particles | $10^{-10}$ [$10^{-12}$ – $10^{-8}$] infections/cell/phage/mL/min |
| $b$ | Burst size: number of phages produced by each infected cell | 50 [5 – 500] phages/cell |
| $\tau$ | Lysis time, the lag between infection of a susceptible cell and lysis to release new phages | 31.6 [10 – 100] minutes |
| $z$ | Scaling parameter for rate of superinfection of infected cells by free phages relative to rate of infection of susceptible cells | 1 |
| $d$ | Fraction of nutrients replenished by each cell death | 0 [0 – 1] |
| $h$ | Transition rate of susceptible cells to resistant cells, relative to growth rate | 0 [0, $10^{-8}$ – $10^{-1}$] |
| $u_R$ | Maximum per-capita growth rate of resistant cells | 0 [0 – 0.0179] /min |
| $f_a$ | Degree of infection rate plasticity | 0 [0 – 3] |
| $S_0$ | Initial density of susceptible cells | $10^6$ [$10^6$ – $10^8$] CFU/mL |
| $I_0$ | Initial density of infected cells | 0 CFU/mL |
| $P_0$ | Initial density of phages | $10^4 [10^4 - 10^6]$ PFU/mL |
| $N_0$ | Initial density of nutrients | $k - S_0 - I_0$ CFU/mL |

This model produces growth curves under the assumption that bacteria remain fully susceptible to phage infection at all times. However, susceptibility to phage infection is known to decline as bacterial growth and metabolism slow as the population enters stationary phase (51, 55–59). We identified simulated growth curves where this assumption is likely violated when *S* + *I* ≥ 0.9 * *k* at any point in time. In some plots in the main text these simulations are excluded (as noted in the figure legends), but plots with all simulations are included in the supplement.

### Models of bacterial population dynamics with plasticity or evolution

To explicitly model changes in bacterial susceptibility over time via plasticity or evolution (Fig 8), we used previously-published approaches (51) to model three scenarios. In the first scenario, as nutrient density and bacterial growth rate decline, phage growth slows via decreases in the infection rate [Eqs. 5 – 9, (60)] or burst size (Appendix 9), or increases in lysis time (Appendix 9). Here, a_t_ declines linearly with decreasing nutrient availability. The slope of that decline, and the total decrease when nutrients have been completely depleted, is controlled by the f parameter.

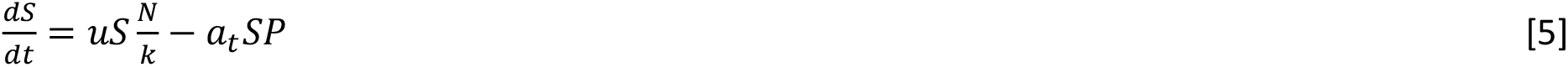

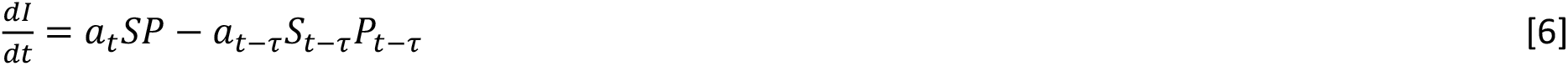

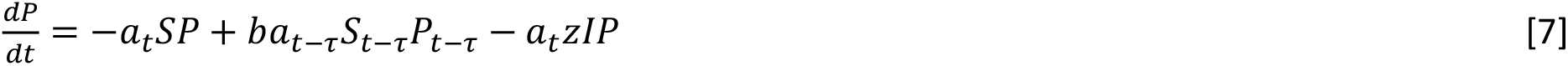

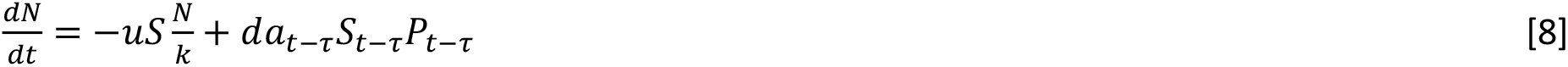

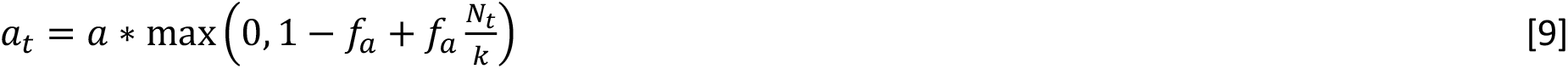

In the second and third scenarios, bacterial cells transition into a resistant state (Eqs. 10 – 12), either because of plastic changes conferring a non-growing resistant state (*u*_*R*_ = 0), or evolutionary mutations conferring cost-free resistance (*u*_*R*_ > 0) [Appendix 9, (61)]. Descriptions of all parameters are provided in Table 1.

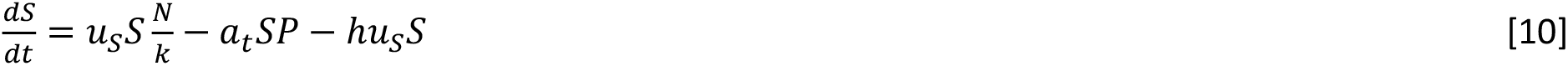

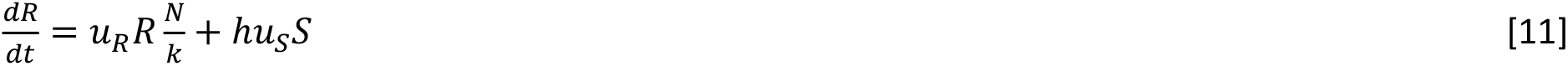

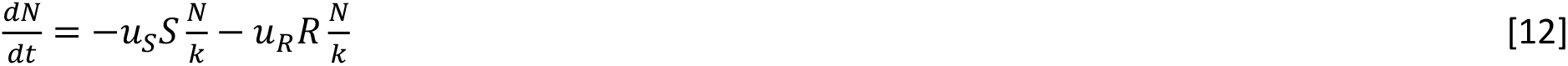

### Simulation implementation and analysis

Simulated population dynamics data were generated using deSolve (62), using default values for all parameters (Table 1) for 48 hours of simulated time unless otherwise noted. Data was analyzed using gcplyr (63): maximum growth rate was calculated using a sliding window 5 data points wide on non-log-transformed data, 10^4^ CFU/mL was used as the arbitrary threshold for extinction time, and 10^6^ CFU/mL was used as the arbitrary threshold for emergence time. Principal component analyses were carried out using each timepoint as a variable, using either the raw density values or (for ‘normalized PCA’) the difference in density values from a control with no phages added. Relative area under the curve (AUC) in Fig 7 was calculated as AUC divided by the AUC of a control curve with no phages added. The coefficient of variation in Fig 7 is the standard deviation divided by the mean. All code used to generate, analyze, and visualize data in R v4.4.3 is available at https://github.com/mikeblazanin/growth-curves.

## Results

First, we tested how changes in phage traits (burst size, lysis time, and infection rate) alter the shape of the curves of bacterial population dynamics. Regardless of the phage trait that varies, we observe that all bacterial populations still display a characteristic exponential phase, before the population density peaks and then rapidly declines due to phage killing (Fig 1). Notably, phage traits do not substantially alter the bacterial population’s density or growth rate during the initial exponential phase, even though our simulations assume cells cease growth immediately upon infection. We and others have observed such a pattern empirically [Appendix 8, (19, 36, 40, 41, 43–47, 49)], which arises because infection remains rare during the exponential phase.

**Figure 1.**
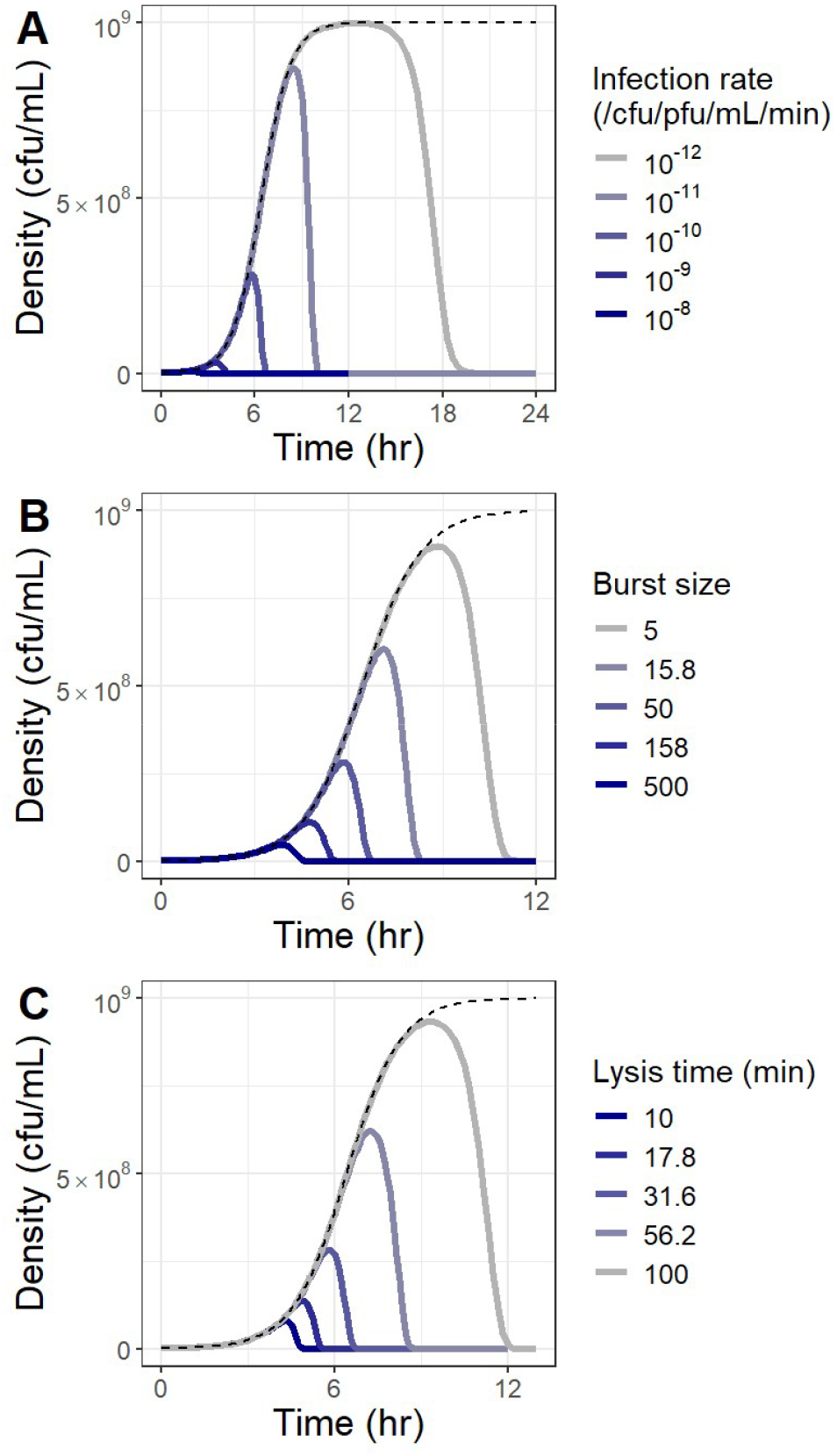
Phage infectivity alters peak and death phases of bacterial population dynamics, but not exponential phase. Bacterial population dynamics were simulated with phages with varying infection rates (A), burst sizes (B), or lysis times (C), plotting the total bacterial density over time. The dashed line denotes bacterial growth in the absence of phages.

Next, we tested how changes in phage traits affect the various metrics that have been proposed (4, 37–46, 48, 50) to quantify phage infectivity from bacterial population dynamics. As expected from the patterns in bacterial population dynamics in Fig 1, metrics like the bacterial growth rate (38, 41), time to reach a threshold density (39, 42), and rate of population decline (41) do not correlate with phage infection rate (Fig 2A, 2B, 2E), burst size (Fig S2), or lysis time (Fig S3). In contrast, metrics like the height and timing of the peak bacterial density (45, 47), the timing of bacterial population extinction (45), the area under the curve (38, 40, 43, 44, 46, 48), and multivariate principal component analyses (37, 48) correlate strongly with phage infection rate (Fig 2C, 2D, 2F, 2G, 2H), burst size (Fig S2), and lysis time (Fig S3).

**Figure 2.**
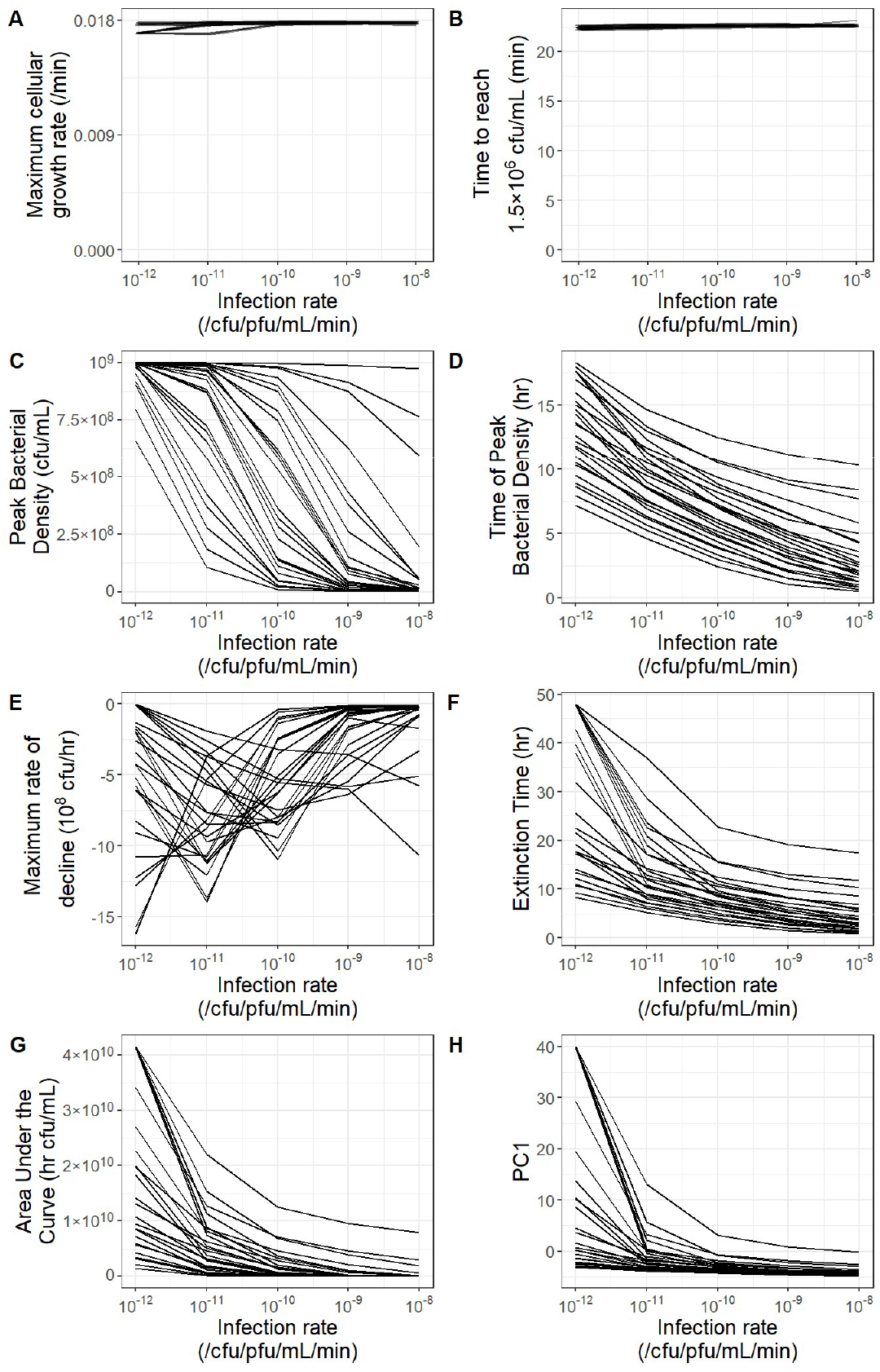
Metrics of the exponential phase and rate of decline do not correlate with phage infectivity, while metrics of peak, overall growth, and timing of death phase do correlate with phage infectivity. Bacterial population dynamics were simulated for 48 hours with phages with all combinations of varying infection rates, lysis times (10, 17.8, 31.6, 56.2, 100 mins), and burst sizes (5, 15.8, 50, 158, 500 PFU/infection). Each line plots the metrics calculated from bacterial population dynamics with phages having the same lysis time and burst size, across varying infection rates. **A, B**. Small amounts of jitter in both the x and y direction were added to aid visualization of many overlapping lines. **F**. Six populations did not reach the extinction threshold and are plotted as 48 hours. **H**. PC1 is the first principal component from a principal component analysis of the bacterial population dynamics.

Next, we tested whether metrics calculated from bacterial population dynamics are correlated with each other, since it remains unclear how these different metrics compare to one another and whether they provide redundant versus complementary information about phage infectivity. We find that different metrics are extremely tightly correlated with one another, suggesting they provide redundant information (Figs 3, S4).

**Figure 3.**
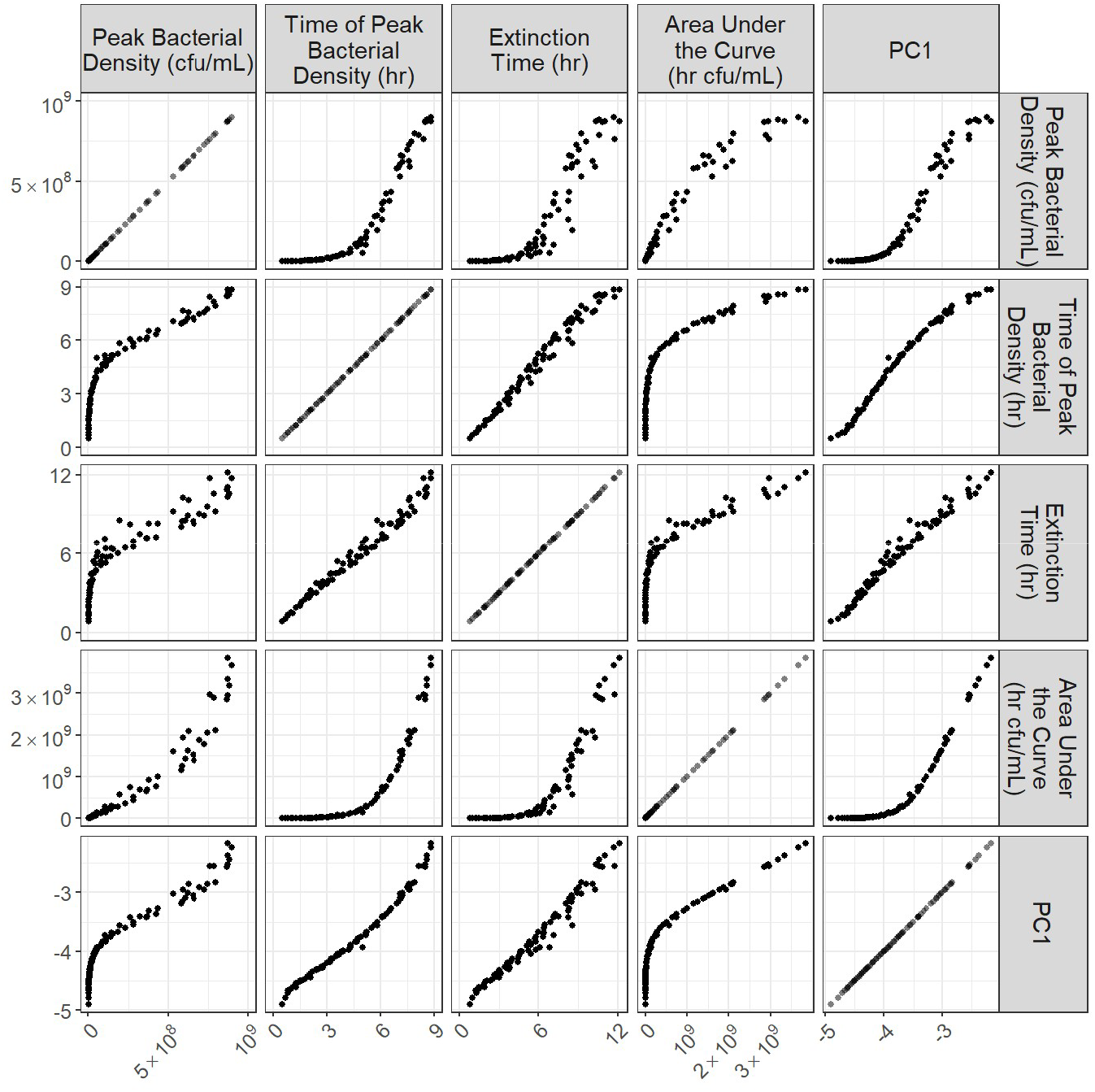
Metrics of bacterial population dynamics are tightly correlated with each other. Bacterial population dynamics were simulated with phages with all combinations of varying infection rates (10^−12^, 10^−11^, 10^−10^, 10^−9^, 10^−8^ /CFU/PFU/min), lysis times (10, 17.8, 31.6, 56.2, 100 mins), and burst sizes (5, 15.8, 50, 158, 500 PFU/infection). Bacterial populations which approximately reached their stationary phase density are excluded from this plot (see Fig S4). PC1 is the first principal component from a principal component analysis of the bacterial population dynamics.

Thus far, we have shown that metrics calculated from bacterial population dynamics correlate strongly with each other and with the phage traits of burst size, lysis time, and infection rate. However, many experiments do not go to the level of quantifying specific phage traits, instead quantifying overall phage growth. Here, we find that metrics calculated from bacterial population dynamics are also strongly correlated with average phage growth rate (Fig 4, Appendix 4). These findings are consistent with past *in vitro* and *in silico* work (45, 47) and suggest that lytic phage growth rate can be inferred simply by quantifying the observable effects the phage has on the population density of its bacterial host.

**Figure 4.**
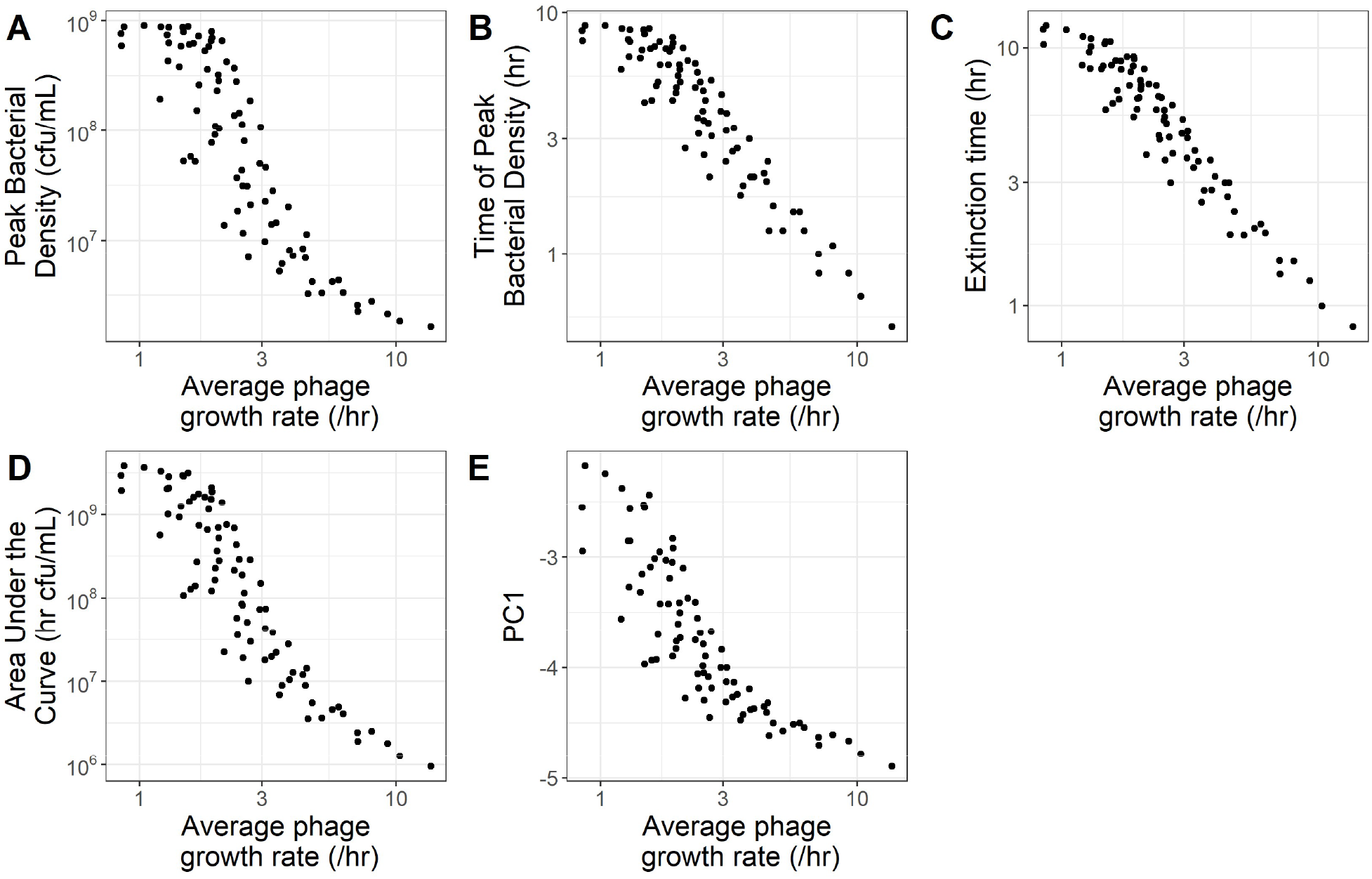
Metrics of bacterial population dynamics are correlated with phage growth rate. Bacterial population dynamics were simulated with phages with all combinations of varying infection rates (10^−12^, 10^−11^, 10^−10^, 10^−9^, 10^−8^ /CFU/PFU/min), lysis times (10, 17.8, 31.6, 56.2, 100 mins), and burst sizes (5, 15.8, 50, 158, 500 PFU/infection). PC1 is the first principal component from a principal component analysis of the bacterial population dynamics. Populations which approximately reached their stationary phase density are not plotted here (see Fig S5). The average phage growth rate was calculated as 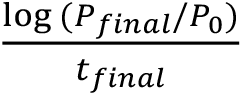 using the extinction time as the final timepoint.

We next quantified how the phage traits of burst size, lysis time, and infection rate interact with each other in altering bacterial population dynamics, as well as quantifying how strongly each trait affects the metrics calculated from bacterial population dynamics. All three traits contribute substantially to shaping bacterial population dynamics (Fig 5, S6 – S10), frequently interacting with diminishing returns (Fig S11). Of the three traits, lysis time typically has the strongest absolute effect (Fig 5D, S12); that is, a given fold-change in lysis time has a larger effect than the same fold-change in burst size or infection rate. However, the three traits are not equally variable: infection rate has the widest range of natural variation, followed by burst size, then lysis time (Table S1). If we normalize to compare across the full range of natural variation, infection rate often has the largest effect (Fig 5E, S12). Regardless, these data show that metrics of bacterial population dynamics provide an effective way to quantify the combined effects of multiple phage traits without measuring individual phage traits.

**Figure 5.**
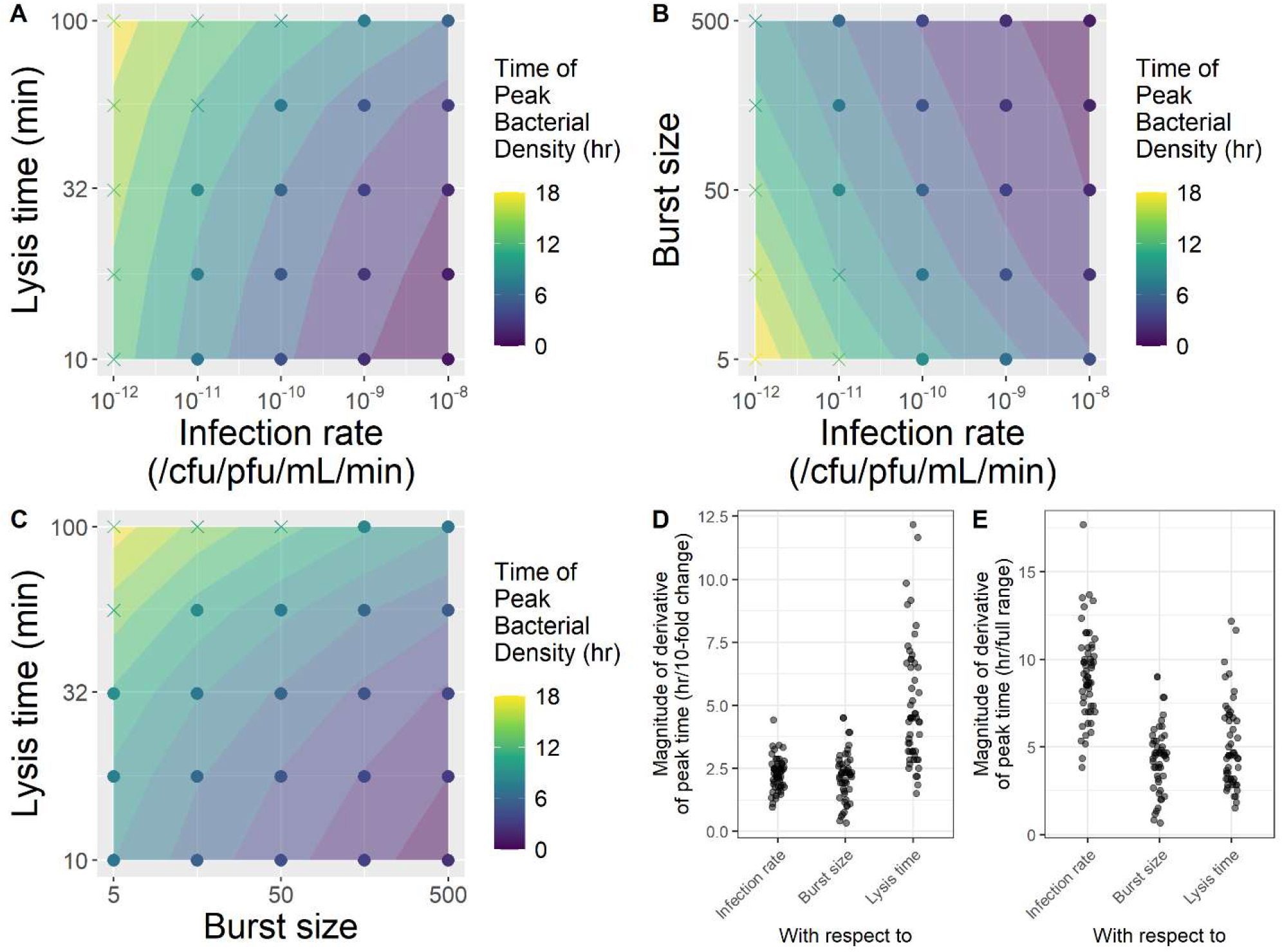
Phage traits jointly determine time of peak bacterial density. Bacterial population dynamics were simulated with phages with all combinations of varying infection rates (10^−12^, 10^−11^, 10^−10^, 10^−9^, 10^−8^ /CFU/PFU/min), lysis times (10, 17.8, 31.6, 56.2, 100 mins), and burst sizes (5, 15.8, 50, 158, 500 PFU/infection). Plotted are subsets of the simulations where **A**, burst size = 50, **B**, lysis time = 31.6 min, or **C**, infection rate = 10^−10^ /cfu/pfu/mL/min. Bacterial populations which approximately reached their stationary phase density are plotted as ‘x’s. **D**. We calculated the magnitude (i.e. absolute value) of the rate of change of the time of peak bacterial density against a 10-fold change in each phage trait. **E**. We calculated the magnitude (i.e. absolute value) of the rate of change of the time of peak bacterial density against each phage trait normalized to have a range of 1. Populations which approximately reached their stationary phase density are not plotted in D and E (see Fig S12).

Next, we tested how strongly the initial densities of bacteria and phages affect the metrics calculated from bacterial population dynamics. Initial densities can substantially alter bacterial population dynamics, with initial bacterial density typically having a stronger effect than initial phage density (Fig 6, Appendix 6, Figs S13 – S15). Fortunately, both initial bacterial density and initial phage density typically have weaker effects on bacterial population dynamics than phage infectivity, so although initial densities do need to be experimentally controlled, random noise in inoculation densities should not obscure differences in infectivity (Appendix 6, Figs S16, S17). Additionally, in contrast to the strong interactions between *traits* in determining population dynamic metrics (Fig 5, Appendix 5), the effects of log initial phage and log bacterial density are approximately linear on many metrics (Fig 6, Appendix 6). Thus, experimenters can manipulate initial bacterial and phage density to maximize the signal of phage infectivity between strains (Fig S14C, S14D).

**Figure 6.**
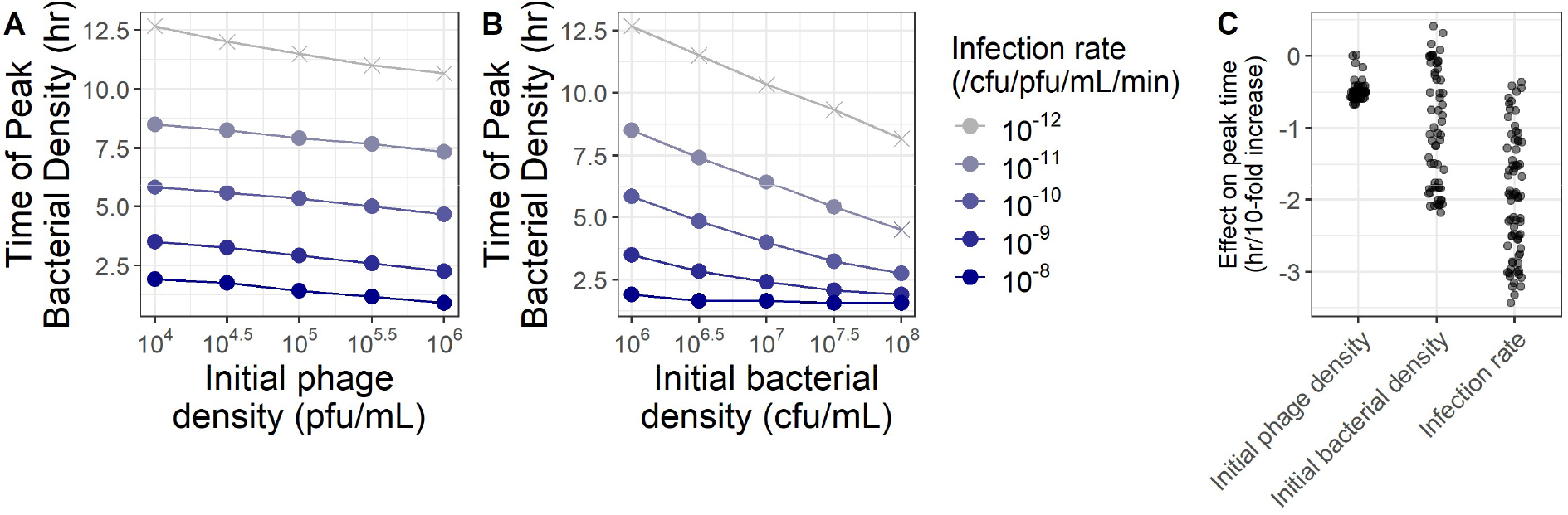
Metrics of bacterial population dynamics can be sensitive to inoculation densities. Bacterial population dynamics were simulated with phages with all combinations of varying infection rates and initial phage (A) or bacterial (B) densities (while holding the other constant). Bacterial populations which approximately reached their stationary phase density are plotted as ‘x’s. **A**. The initial density of bacteria was held constant at 10^6^ CFU/mL. **B**. The initial density of phages was held constant at 10^4^ PFU/mL. **C**. We calculated the magnitude of the rate of change of the time of peak bacterial density against a 10-fold change in infection rate, initial bacterial density, or initial phage density. Populations which approximately reached their stationary phase density are not plotted (see Fig S15).

We now set out to test how metrics of population dynamics can be used to compare phage infectivity across different bacterial hosts, for instance when bacterial strains vary in their resistance to infection. Many population dynamic metrics used to infer infectivity can be strongly affected by variation in bacterial traits like growth rate or stationary phase density (Fig 7, Appendix 7). Of the metrics, the time of peak bacterial density and extinction time tend to be the best metrics (i.e., are least affected by variation in bacterial traits). One proposed approach to explicitly account for variation in bacterial traits is to calculate the area under the curve (AUC) relative to the AUC of a control where bacteria are grown alone (40, 44, 46). When bacterial strains vary in their growth rate, relative AUC can be a somewhat better indicator than raw AUC of infectivity (Fig 7A). However, counterintuitively, when bacterial strains vary in their stationary phase density, relative AUC is actually a much worse indicator of infectivity (Fig 7B). This arises because phages typically cause bacterial populations to collapse before they begin to approach stationary phase, so AUC values vary more in the control than in the presence of phages. A second approach to explicitly account for variation in bacterial traits is to use multivariate ordination methods like PCA on the raw density values or the density values relative to those of a control where bacteria are grown alone (37, 48). When bacteria vary in growth rate, both PCA approaches work well as metrics of infectivity, with little overall improvement from PCA on relative densities (Fig 7A). However, when bacterial strains vary in their stationary phase density, PCA on relative densities is a much worse metric of phage infectivity (Fig 7B). This arises for much the same reason as AUC: when bacteria vary in stationary phase density, control densities vary more than densities in the presence of phages.

**Figure 7.**
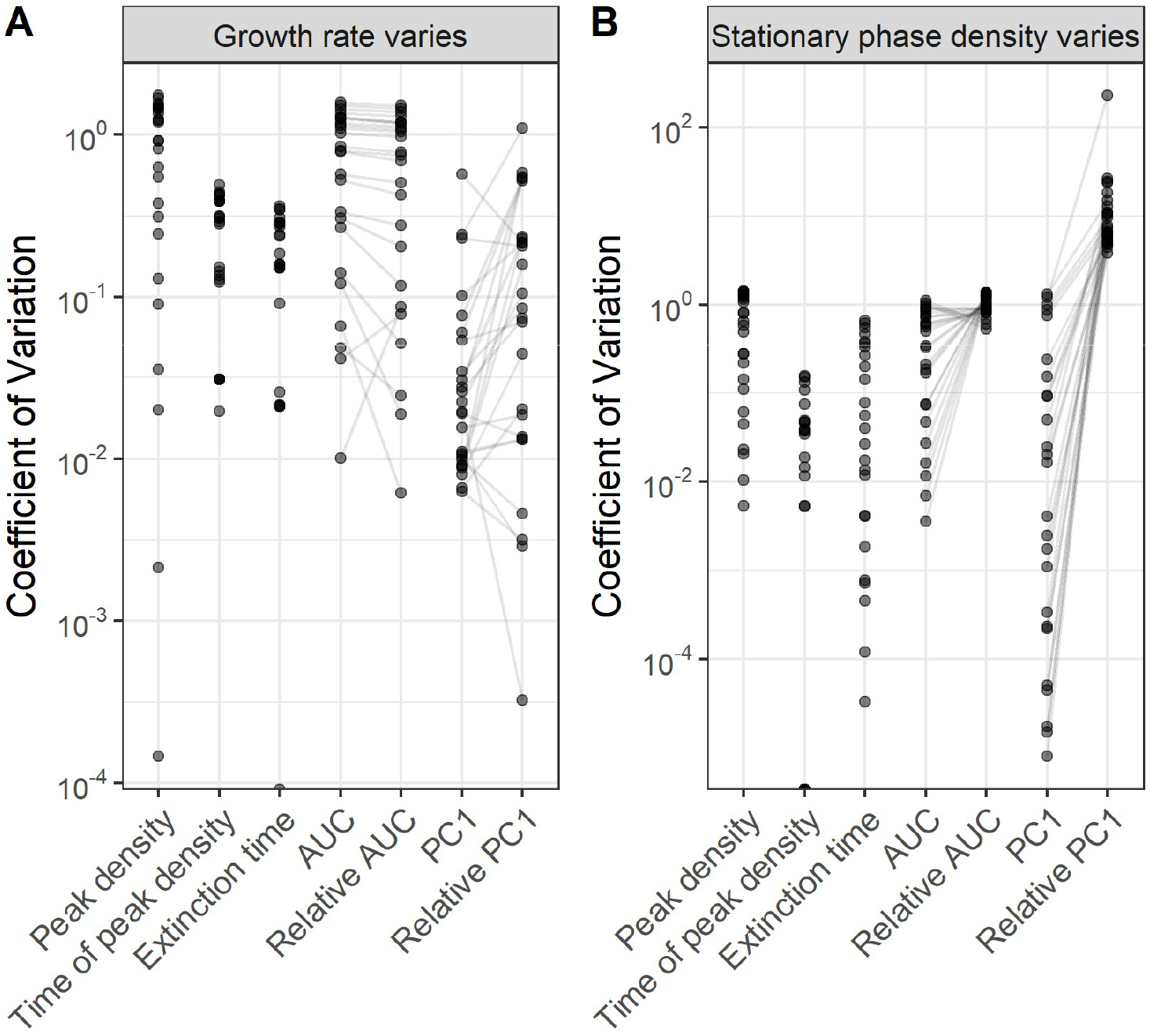
When bacteria vary the time of peak bacterial density and extinction time are the best metrics of phage infectivity. Bacterial population dynamics were simulated with phages with all combinations of varying infection rates (10^−12^, 10^−11^, 10^−10^, 10^−9^, 10^−8^ /CFU/PFU/min) and bacteria with varying stationary phase densities (10^8^, 10^8.5^, 10^9^, 10^9.5^, 10^10^ CFU/mL) and growth rates (0.04, 0.027, 0.018, 0.012, 0.008 /min; doubling times of 17, 26, 39, 58, and 87 mins). We then calculated the amount of variation in each metric among simulations with **A**. the same infection rate and stationary phase density, or **B**. the same infection rate and growth rate. Smaller coefficients of variation indicate that the measure is a better indicator of infectivity across bacteria that vary in **A**. growth rate, or **B**. stationary phase density. For AUC and Relative AUC, and PC1 and Relative PC1, lines connect sets of simulations with the same parameter values.

Finally, we sought to test how bacterial population dynamics can be used to quantify the effects of phages over longer timescales, when bacteria can become resistant through plastic or evolutionary changes. For instance, *in vitro* bacterial susceptibility is often observed to decline as bacterial growth slows (51, 55–59), and bacteria are also known to readily evolve resistance against phages. To assess these effects on bacterial population dynamics, we used previously-published approaches (51) to simulate three scenarios (Fig 8, Appendix 9):

**Figure 8.**
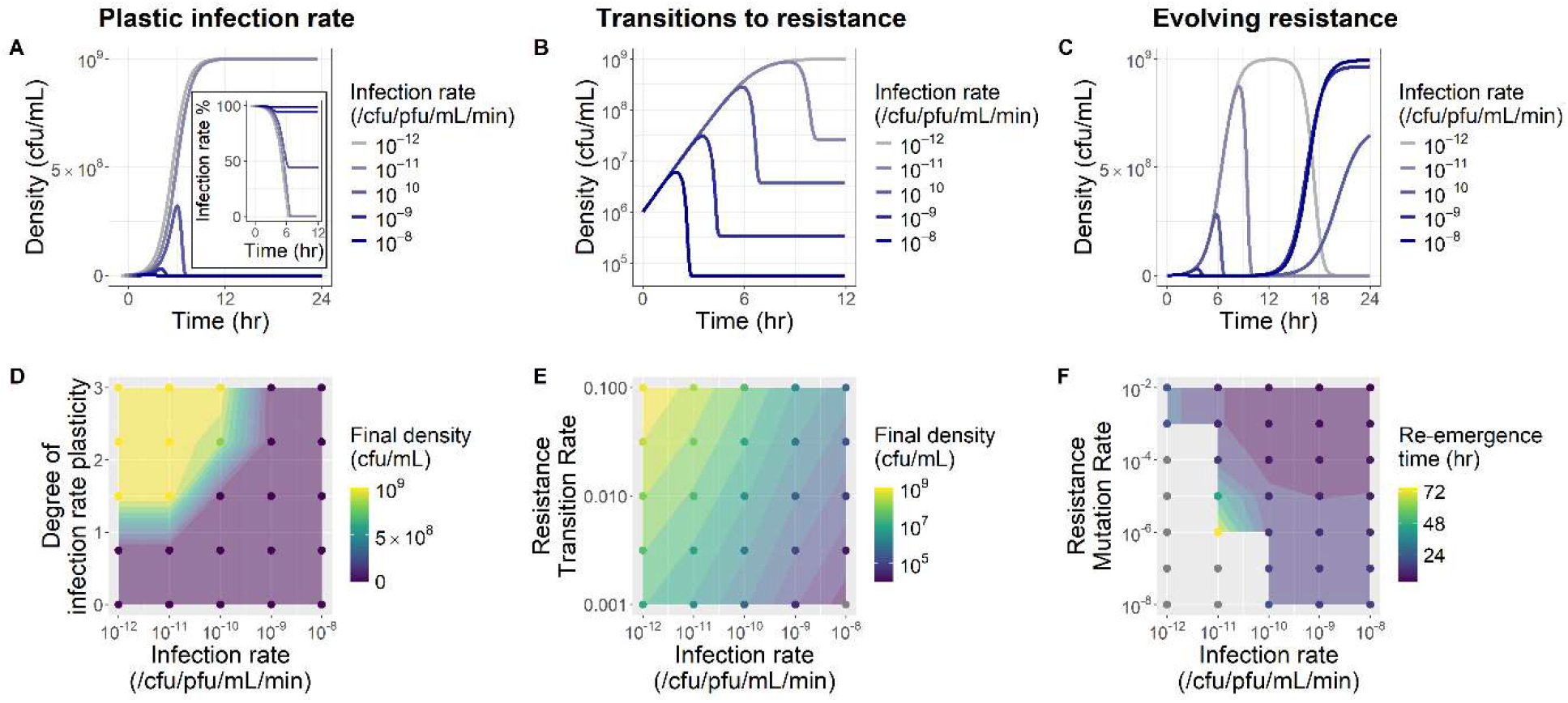
Plasticity and evolution can alter the shape of bacterial population dynamics over longer timescales. Bacterial population dynamics were simulated with phages with varying infection rates (10^−12^, 10^−11^, 10^−10^, 10^−9^, 10^−8^ /CFU/PFU/min). **A**. Phages were simulated with an infection rate that declines linearly with declining bacterial growth rate (f_a_ = 1.5). Bacterial populations that reach stationary phase enough that the phage infection rate reaches 0 (inset) can persist without ever crashing. Population densities and infection rates have been slightly offset horizontally for visualization. **B**. Bacteria were simulated with transitions into a non-growing resistant state (*h* = 0.01). **C**. Bacteria were simulated with cost-free mutations providing complete phage resistance (*h* = 10^−5^).

1. Phage growth weakening as bacterial growth slows (20, 51, 57, 60, 64, 65). Here, we find that bacterial population dynamics are substantially altered if bacteria reach stationary phase (Fig 8A). There, bacterial growth slows enough that phage growth falls to zero, preventing the collapse observed in previous simulations. This also makes it impossible to distinguish phage infectivity differences between any bacterial populations that are sufficiently resistant to reach stationary phase (Fig 8D).
2. Bacterial cells transitioning into a phenotypically resistant state (61). Here, transitions into a phenotypically resistant state can produce patterns of partial population collapse (Figs 8B, 8E) that have been observed empirically [(9), Fig S21], although such patterns in optical density can also be explained by debris (Fig S27).
3. Bacteria evolving mutations that confer resistance to phage infection. Here, bacterial populations collapse to near-zero densities before rebounding because of the evolution of phage-resistant mutants (Fig 8C).

Across all three scenarios, metrics like final density or re-emergence time reflect both infectivity and the degree/rate of plasticity or evolution (Fig 8D, 8E, 8F), while metrics like peak density, time of peak density, and extinction time remain good indicators of phage infectivity alone, independent of the effects of plasticity or evolution (Fig S29).

## Discussion

Here, we set out to test how bacterial population dynamics can be used to quantify phage infectivity (and bacterial resistance), using mathematical models to simulate population dynamics with known trait values. We showed that many, but not all, metrics of bacterial population dynamics reflect phage infectivity (Fig 2), and that metrics are strongly correlated with one another (Fig 3). We then showed that these metrics can be used to infer phage growth rate (Fig 4), providing an effective way to quantify the combined effects of multiple phage traits (Fig 5). We also showed that metrics can be somewhat affected by initial inoculum densities (Fig 6), and identified time of peak density and extinction time as the best metrics to compare across different bacterial hosts (Fig 7). Finally, we showed that the effects of phages can sometimes be inferred over longer time-scales where bacterial plasticity or evolution can alter population dynamics (Fig 8).

Observation of bacterial population dynamics complements existing methods for estimating phage infectivity. Existing methods often exhibit tradeoffs between throughput and precision (3, 4), with some approaches (e.g. efficiency of plaquing) providing quantitative but low-throughput measures of infectivity (5–7), while others (e.g. cross-streaks) provide qualitative but high-throughput measures of infectivity (8). In contrast, bacterial population dynamics can be easily scaled to collect many replicates in parallel and can produce quantitative measures of infectivity, albeit with a smaller range of detection (~2 orders of magnitude in infectivity, Fig 2C). In addition, inferring phage infectivity from bacterial population dynamics may better reflect phage-bacteria interactions in liquid environments than the existing agar surface-based methods for observing phage infectivity, although further work is needed.

Our work builds on prior papers that used bacterial population dynamics to infer phage activity (37–50). However, our findings contrast with some of their previously-reported results. For example, several papers have suggested using normalized area under the curve (40, 44, 46) or PCA (37) to compare infectivity across bacterial hosts, but we find that these metrics are not particularly well-suited to this task, and in some cases are worse than the unnormalized metrics (Fig 7). Additionally, two recent papers have suggested fitting mathematical models to bacterial population dynamics to extract phage trait values (47, 49). Although we did not directly explore fitting-based approaches, so they remain an avenue for future theoretical work, our findings that metrics strongly covary (Fig 3) and that many combinations of trait values can produce similar curves (Fig 5) would suggest that bacterial population dynamics are unlikely to be sufficient to quantify specific phage life history traits like adsorption rate, burst size, or lysis time. At the same time, our findings do align with some previously-reported results. For instance, we found that the exponential phase of population dynamics provides little information about phage infectivity (Fig 1), a pattern that has been observed in our own empirical data (Appendix 8), previous phage-bacteria studies (19, 36, 40, 41, 43–47, 49), and in epidemiology (66, 67). Our simulations also reproduced previously-reported near-linear relationships between phage growth rate and bacterial extinction time [Fig 4C, (45)], peak bacterial density and initial phage density [Fig S14B, (47)], and time of peak bacterial density and initial bacterial density [Fig 6B, (49)].

Our work has established a broad foundation for using bacterial population dynamics to quantify phage infectivity and bacterial resistance, opening avenues for future work. In particular, empirical work is needed to test the patterns and predictions from this paper. This includes experimental validation of the relationships among metrics (Fig 2) and between metrics and phage growth rate (Fig 4), but especially to quantify relationships between phage traits and population dynamic metrics (Figs 3, 5, S2, S3).

Additional empirical work is also needed to strengthen our understanding of the relationship between cell density and measured proxies of cell density like optical density (68–71), especially over longer timescales where some cells may become resistant or debris may play a role [Figs 8, S27, (9)]. Both empirical and theoretical work are needed to better understand how bacterial susceptibility to phages changes across the phases of bacterial growth (51, 55–59), something our simulations generally ignored (but see Fig 8). At the same time, future theoretical work should test the capacity and limitations of fitting-based approaches to quantify phage infectivity from bacterial population dynamics. Theory should also explore how additional biological processes alter population dynamics and the inference of infectivity, including lysogeny, stochasticity, failed infections, coinfection exclusion, cooperation, or continuous intrapopulation trait variation. Finally, theory should also be applied to improve our understanding of other methods of quantifying infectivity, like the efficiency of plaquing assay (72).

In all, we have shown that bacterial population dynamics can enable powerful quantification of phage-bacteria interactions. Given that such approaches are already widely used heuristically in the phage-bacteria literature, our findings suggest that they may be ripe for quantitative application from basic to applied questions.

## Acknowledgements

Thanks to Alita Burmeister, Tiffany Hamidjaja, Catherine Hernandez, Jordan Lewis, Albert Vill, and Caroline Turner for feedback on the manuscript. Thanks to members of the Turner lab for feedback on the work in this project. Thanks to Teresa Carter for contributions to very early work on this project. Thanks to the Yale Institute for Biospheric Studies and Howard Hughes Medical Institute (HHMI) for funding supporting this work. Thanks to the Yale STARS program for funding supporting WA during this project.

