## Supplementary Material for "Quantifying phage infectivity from characteristics of bacterial population dynamics"

### Appendices

#### Appendix 1. Modeling approach

A.
$$\frac{dS}{dt} = uS\frac{N}{k} - aSP$$
$$\frac{dI}{dt} = aSP - aS_{t-\tau}P_{t-\tau}$$
$$\frac{dP}{dt} = -aSP + baS_{t-\tau}P_{t-\tau} - azIP$$
$$\frac{dN}{dt} = -uS\frac{N}{k} + daS_{t-\tau}P_{t-\tau}$$

B. C.

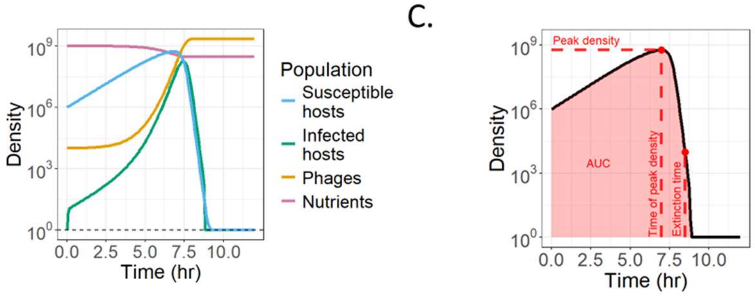


**Figure S1. Conceptual figure showing how we simulated bacterial population dynamics with phages. A.** A modified SI-type system of delay differential equations was used to model the growth of bacterial and phage populations over time. Susceptible bacteria (*S*) grow logistically (at rate *u*) up to a stationary phase density (*k*). Phage virions (*P*) infect susceptible hosts (at a rate *a*) and after a discrete lysis time ($\tau$) release progeny virus particles (*b* per infection). Phages can also superinfect cells (at a rate *z*, relative to *a*). Lysis of cells releases nutrients (*N*) back into the environment (at fraction *d*). **B.** Simulation of this model (with parameters $a = 10^{-10}$ /cfu/pfu/mL/min, $b = 5$, and $\tau = 10$ mins) shows initial growth of phage population and susceptible and infected bacterial populations, before bacteria go extinct and phage growth ceases. **C.** In practice in the laboratory, the typical available data is the total bacterial population ($S + I$), plotted in black. Growth curve metrics can then be extracted. Shown here are peak density and time, time when density drops below a threshold (‘extinction’ time), and area under the curve (AUC).

**Table S1.** Experimental studies were surveyed to identify appropriate ranges of values to use for parameters in our simulations.

| Source | Species (Bacteria – Phage) | Adsorption rate (/PFU/CFU/mL/min) | Lysis time (min) | Burst size | Resistance mutation rate |
|---|---|---|---|---|---|
| (73) | *E. coli* – lambda | $1.3 - 9.9 \times 10^{-9}$ | 29.3 – 68 | - | - |
| (74) | *E. coli* – T7 | $3.3 - 4 \times 10^{-9}$ | 10.4 – 12.4 | 266 – 327 | - |
| (75) | *E. coli* – lambda | - | 60 | 4.5 – 620 | - |
| (32) | *E. coli* – lambda | $2.8 \times 10^{-12}$ | 28.3 – 63 | 9.7 – 255 | - |
| (76) | *E. coli* – lambda | $7.5 \times 10^{-11} - 10^{-8}$ | | < 1000 | - |
| (77) | *E. coli* – T4 | - | 37.5 – 53 | - | - |
| (45) | *P. syringae* – Phi6 | $10^{-12} - 10^{-9}$ | 90 – 105 | 100 – 160 | - |
| (78) | *P. aeruginosa* – various | $2.8 \times 10^{-12} - 1.7 \times 10^{-11}$ | 45 – 100 | 37 – 589 | - |
| (79) | Various | $5.5 \times 10^{-11} - 3.7 \times 10^{-9}$ | 13 – 60 | 50 – 3570 | - |
| (80) | *E. coli* – lambda | - | 52.5 – 103.3 | 104 – 186 | - |
| (81) | *E. coli* – various | $5 \times 10^{-9} - 1.2 \times 10^{-8}$ | 9 – 19 | 161 – 345 | - |
| (82) | *P. aeruginosa* – various | - | - | - | $10^{-7} - 10^{-2}$ |
| (83) | *E. coli* – U136B | - | - | - | $4 \times 10^{-6}$ |
| (84) | *E. faecalis* - various | - | - | - | $10^{-7} - 10^{-5}$ |
| (85) | *E. coli* – various | - | - | - | $10^{-8} - 10^{-5}$ |
| (86) | *S. cremoris* – various | - | - | - | $10^{-7} - 10^{-4}$ |

### Appendix 2. Correlations between metrics and phage traits

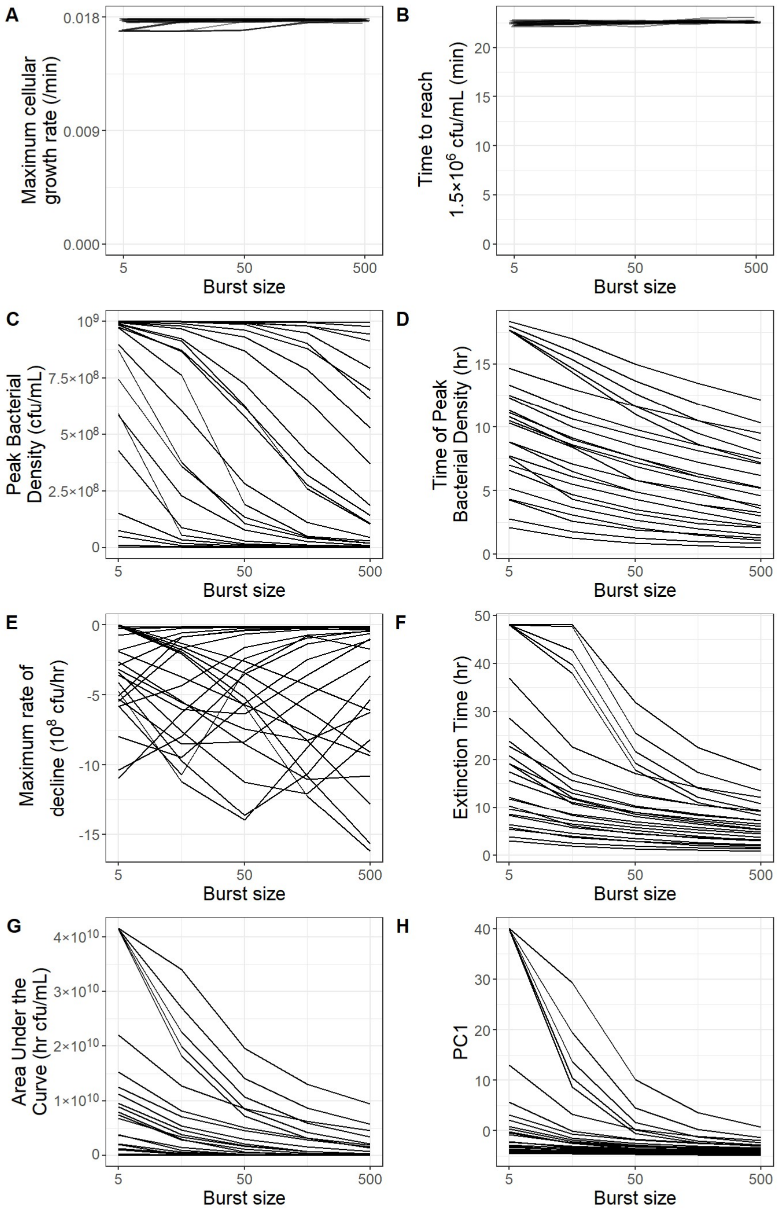


**Figure S2. Metrics of the exponential phase and rate of decline do not correlate with phage burst size, while metrics of peak, overall growth, and timing of death phase do correlate with phage burst size**. Bacterial population dynamics were simulated with phages with varying infection rates ($10^{-12}$, $10^{-11}$, $10^{-10}$, $10^{-9}$, $10^{-8}$ /CFU/PFU/min), lysis times (10, 17.8, 31.6, 56.2, 100 mins), and burst sizes. All other parameters were default values (Table 1). Each line plots the metrics calculated from bacterial population dynamics with phages having the same infection rate and lysis time, across varying burst sizes. In F, $10^4$ CFU/mL was used as the arbitrary threshold for extinction, which six populations did not reach within 48 hours. Small amounts of jitter in both the x and y direction were added to A and B to aid visualization of many overlapping lines.

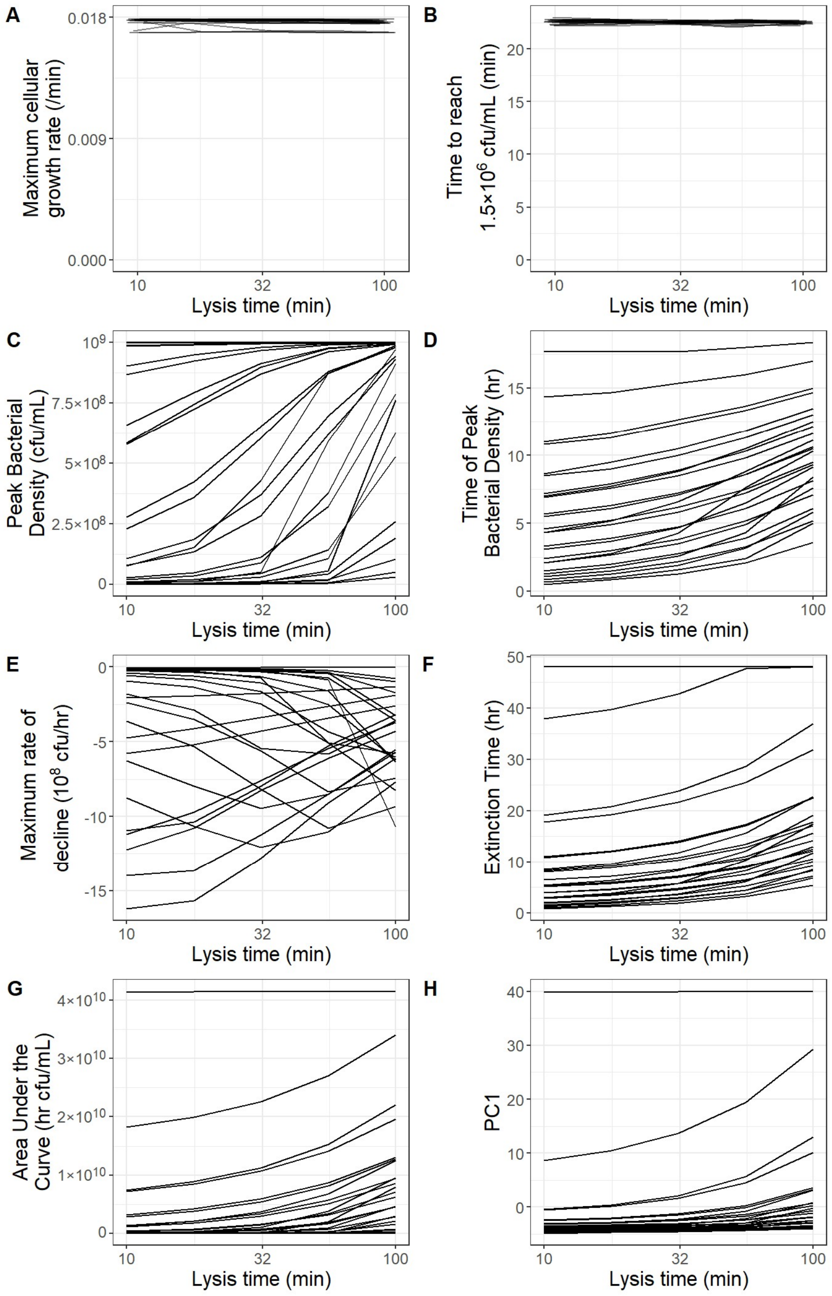


**Figure S3. Metrics of the exponential phase and rate of decline do not correlate with phage lysis time, while metrics of peak, overall growth, and timing of death phase do correlate with phage lysis time**. Bacterial population dynamics were simulated with phages with varying infection rates ($10^{-12}$, $10^{-11}$, $10^{-10}$, $10^{-9}$, $10^{-8}$ /CFU/PFU/min), lysis times, and burst sizes (5, 15.8, 50, 158, 500 PFU/infection). All other parameters were default values (Table 1). Each line plots the metrics calculated from bacterial population dynamics with phages having the same infection rate and burst size, across varying lysis times. In F, $10^4$ CFU/mL was used as the arbitrary threshold for extinction, which six populations did not reach within 48 hours. Small amounts of jitter in both the x and y direction were added to A and B to aid visualization of many overlapping lines.

#### Appendix 3. Relationships between metrics

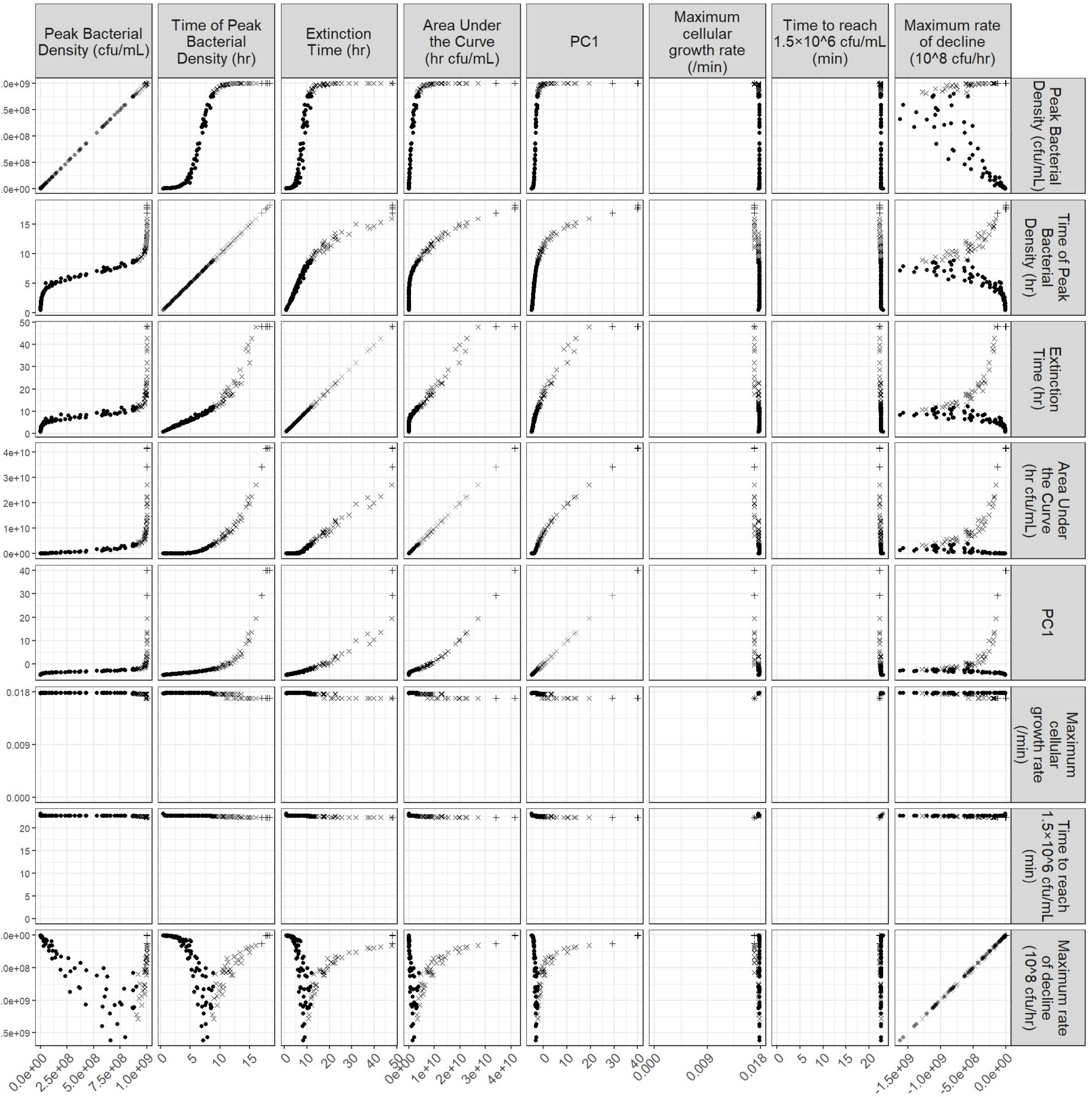


**Figure S4. Metrics of bacterial population dynamics are tightly correlated with each other.** Bacterial population dynamics were simulated with phages with varying infection rates ($10^{-12}$, $10^{-11}$, $10^{-10}$, $10^{-9}$, $10^{-8}$ /CFU/PFU/min), lysis times (10, 17.8, 31.6, 56.2, 100 mins), and burst sizes (5, 15.8, 50, 158, 500 PFU/infection). All other parameters were default values (Table 1). $10^4$ CFU/mL was used as the arbitrary threshold to calculate extinction time. Bacterial populations which approximately reached their stationary phase density are plotted as ‘x’s, and bacterial populations that did not reach extinction within 48 hours are plotted as ‘+’s. PC1 is the first principal component from a principal component analysis of the bacterial population dynamics.

#### Appendix 4. Relationships between metrics and phage growth rate

Past work has shown that the average phage growth rate during a growth curve has a negative relationship with bacterial extinction time both *in vitro* and *in silico* (45). We simulated growth curves using phages with a wide array of trait values, observing that our model reproduces this pattern (Fig S5C).

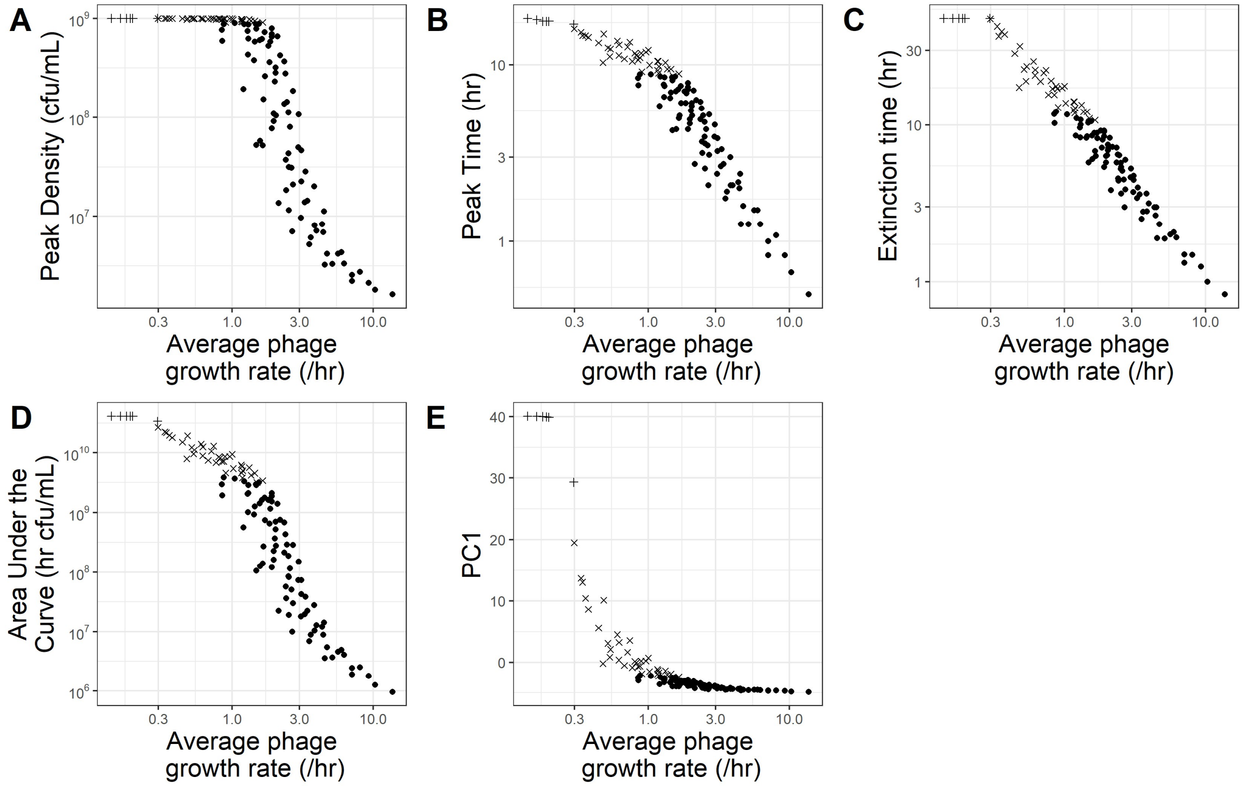


**Figure S5. Metrics of bacterial population dynamics are correlated with phage growth rate.** Bacterial population dynamics were simulated with phages with varying infection rates ($10^{-12}$, $10^{-11}$, $10^{-10}$, $10^{-9}$, $10^{-8}$ /CFU/PFU/min), lysis times (10, 17.8, 31.6, 56.2, 100 mins), and burst sizes (5, 15.8, 50, 158, 500 PFU/infection). All other parameters were default values (Table 1). $10^4$ CFU/mL was used as the arbitrary threshold to calculate extinction time. Bacterial populations which approximately reached their stationary phase density are plotted as ‘x’s, and bacterial populations that did not reach extinction within 48 hours are plotted as ‘+’s. PC1 is the first principal component from a principal component analysis of the bacterial population dynamics. The average phage growth rate was calculated as $\frac{\log(P_{final}/P_0)}{t_{final}}$ using the extinction time as the final timepoint.

#### Appendix 5. The interacting effects of phage trait values on metrics calculated from bacterial population dynamics

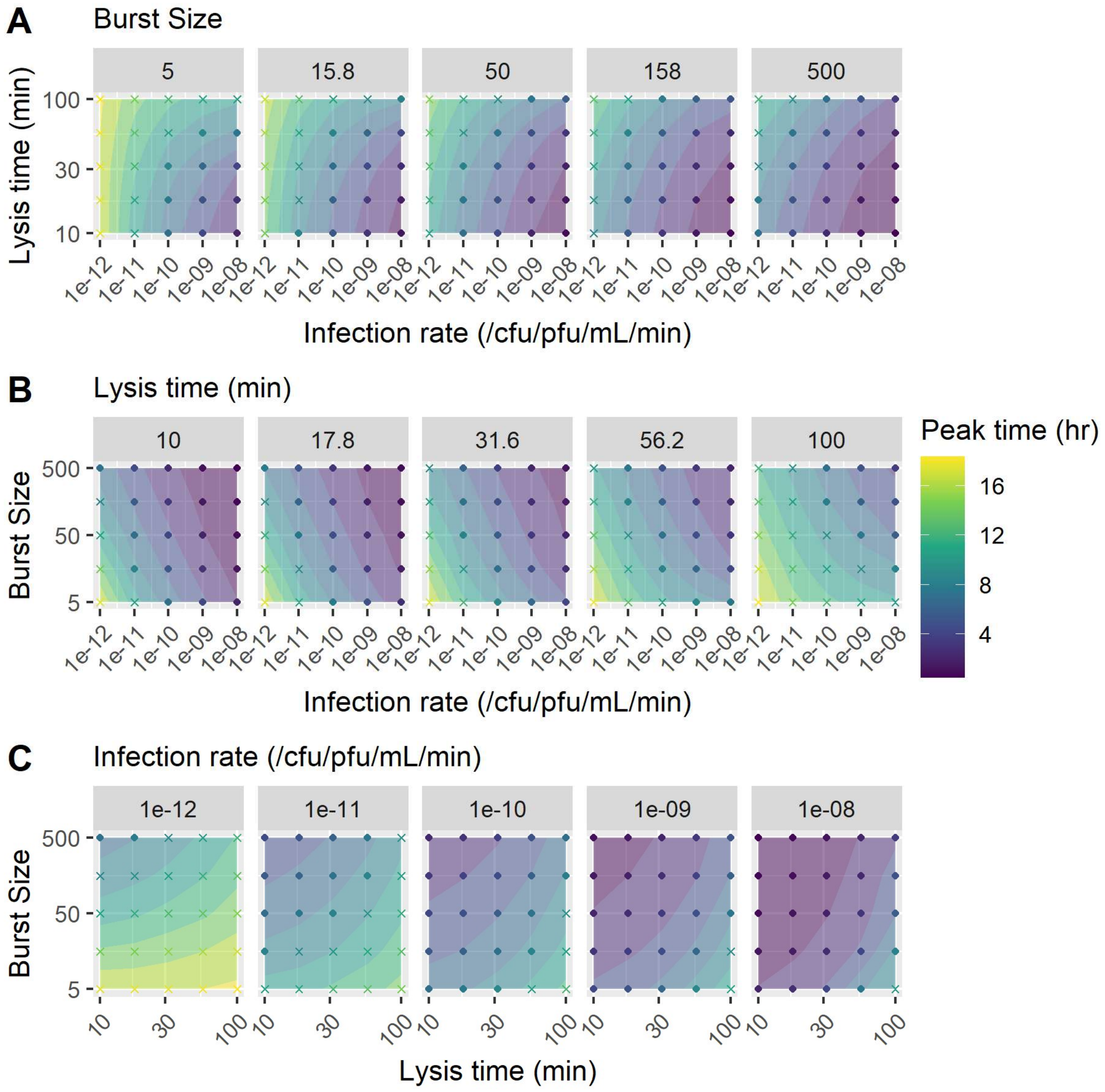


**Figure S6. Phage traits jointly determine time of peak bacterial density.** Bacterial population dynamics were simulated with phages with varying infection rates, lysis times, and burst sizes. All other parameters were default values (Table 1). Bacterial populations which approximately reached their stationary phase density are plotted as 'x's.

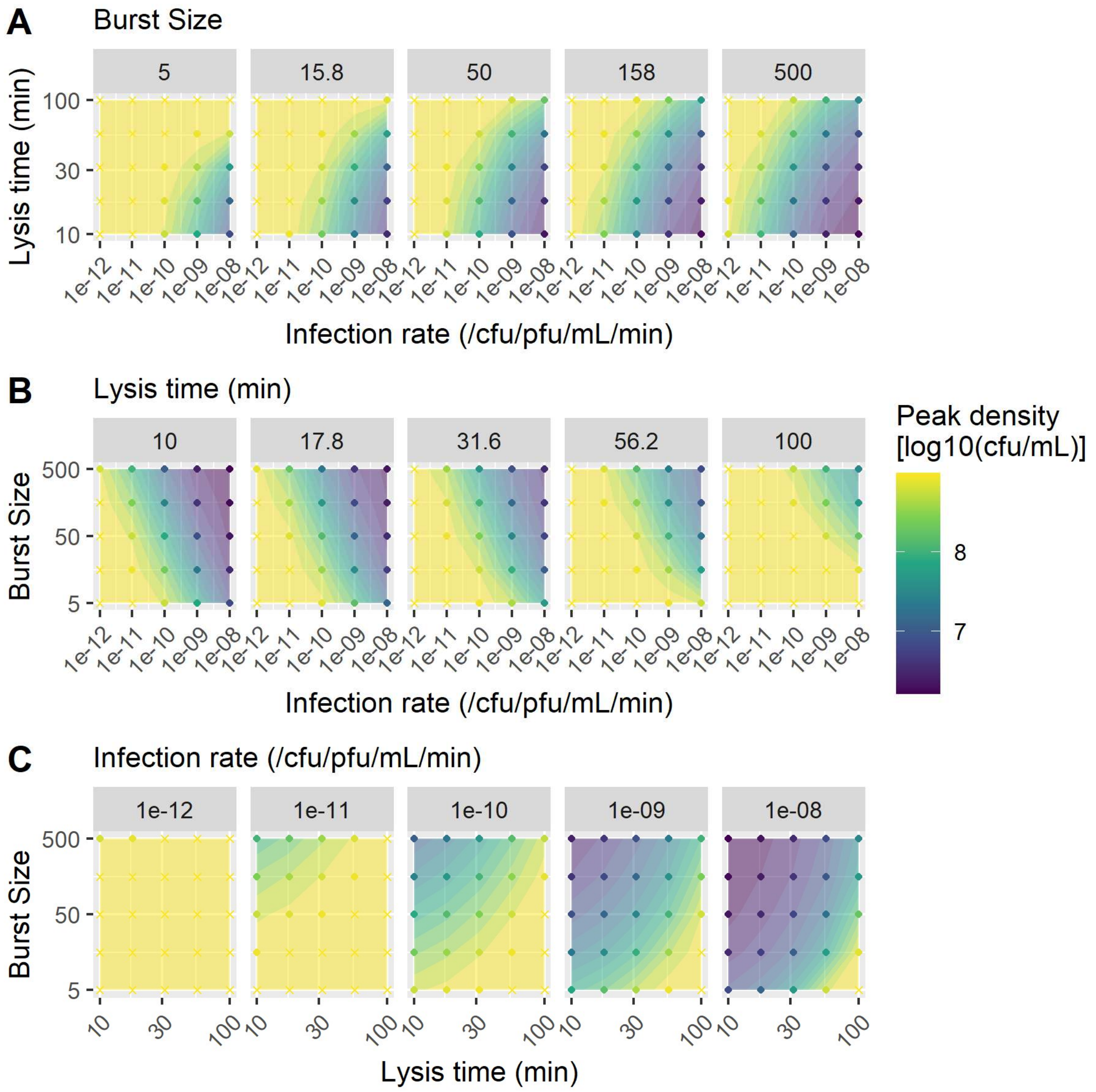


**Figure S7. Phage traits jointly determine peak bacterial density.** Bacterial population dynamics were simulated with phages with varying infection rates, lysis times, and burst sizes. All other parameters were default values (Table 1). Bacterial populations which approximately reached their stationary phase density are plotted as 'x's.

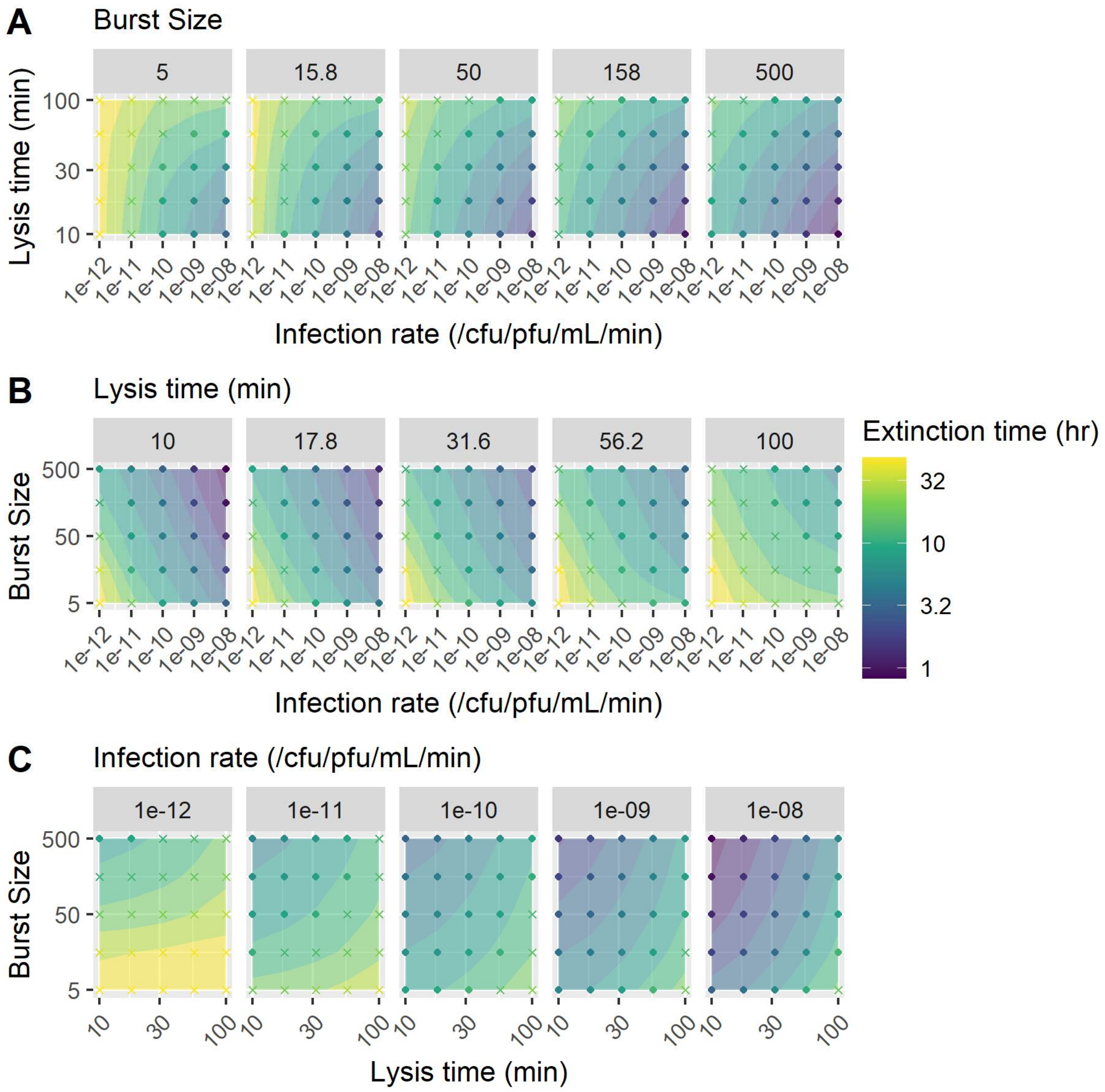


**Figure S8. Phage traits jointly determine bacterial extinction time.** Bacterial population dynamics were simulated with phages with varying infection rates, lysis times, and burst sizes. All other parameters were default values (Table 1). Bacterial populations which approximately reached their stationary phase density are plotted as 'x's.

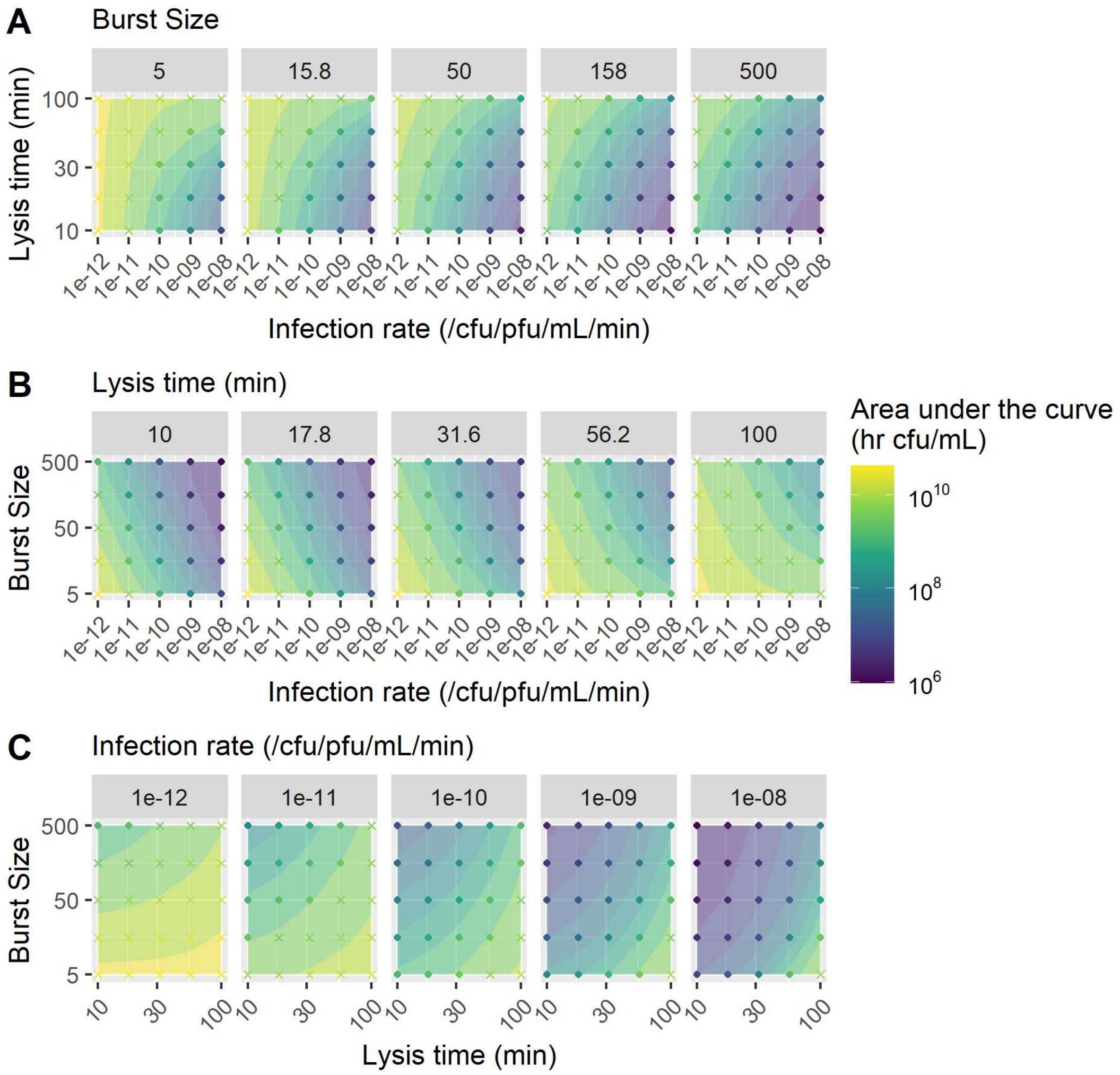


**Figure S9. Phage traits jointly determine bacterial area under the curve.** Bacterial population dynamics were simulated with phages with varying infection rates, lysis times, and burst sizes. All other parameters were default values (Table 1). Bacterial populations which approximately reached their stationary phase density are plotted as ‘x’s.

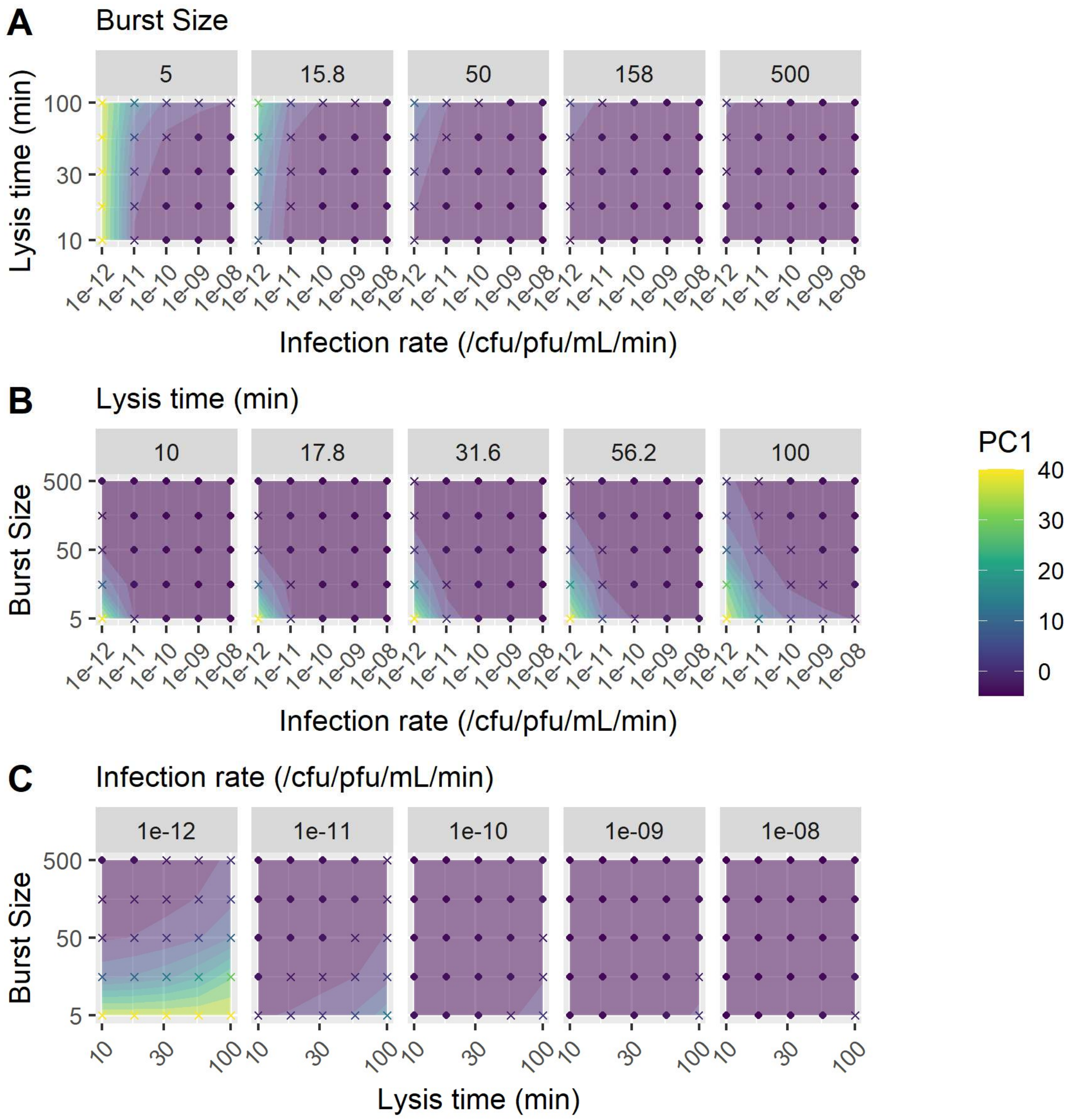


**Figure S10. Phage traits jointly determine the first principal component from a principal component analysis of the bacterial population dynamics.** Bacterial population dynamics were simulated with phages with varying infection rates, lysis times, and burst sizes. All other parameters were default values (Table 1). Bacterial populations which approximately reached their stationary phase density are plotted as 'x's.

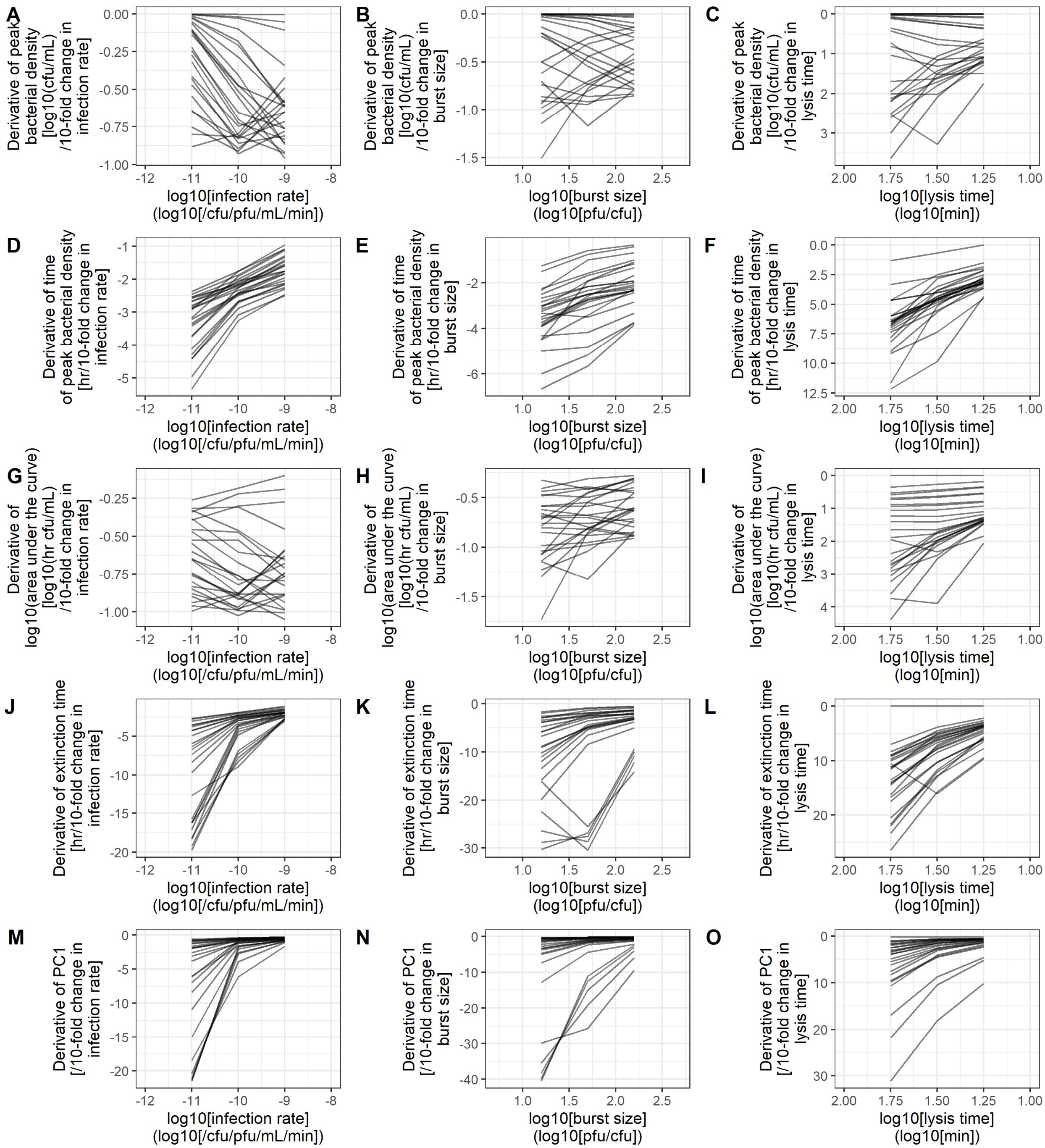


**Figure S11. Phage traits frequently interact with diminishing returns to determine metrics of bacterial population dynamics.** Bacterial population dynamics were simulated with phages with varying infection rates ($10^{-12}$, $10^{-11}$, $10^{-10}$, $10^{-9}$, $10^{-8}$ /CFU/PFU/min), lysis times (10, 17.8, 31.6, 56.2, 100 mins), and burst sizes (5, 15.8, 50, 158, 500 PFU/infection). All other parameters were default values (Table 1). We calculated the rate of change of each metric against a 10-fold change in each phage trait. Each line plots the metrics calculated from bacterial population dynamics with phages having the same values of the other two phage traits.

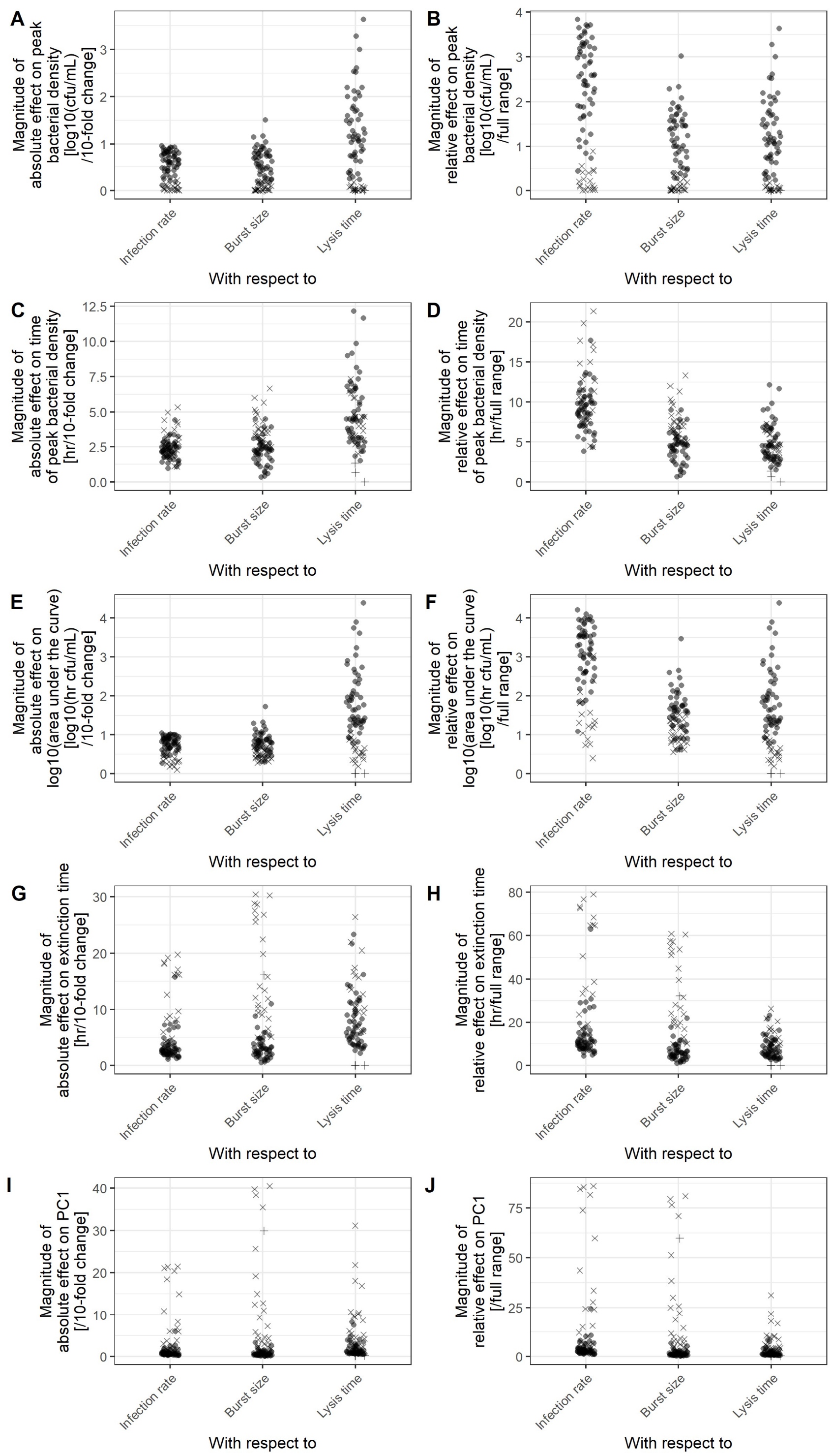


**Figure S12. Lysis time is often the phage trait which most strongly affects metrics of bacterial population dynamics absolutely, while infection rate is often the trait which most strongly affects metrics given the range of natural variation.** Bacterial population dynamics were simulated with phages with varying infection rates ($10^{-12}$, $10^{-11}$, $10^{-10}$, $10^{-9}$, $10^{-8}$ /CFU/PFU/min), lysis times (10, 17.8, 31.6, 56.2, 100 mins), and burst sizes (5, 15.8, 50, 158, 500 PFU/infection). All other parameters were default values

(Table 1). $10^4$ CFU/mL was used as the arbitrary threshold to calculate extinction time. **A,C,E,G,I.** We calculated the magnitude (i.e. absolute value) of the rate of change of each metric against a 10-fold change in each phage trait. Here, we see that a 10-fold change in lysis time often has a larger effect than the same fold-change in burst size or infection rate. **B,D,F,H,J.** Since infection rate, burst size, and lysis time are not equally variable, we calculated the magnitude (i.e. absolute value) of the rate of change of each metric against each phage trait normalized to have a range of 1. Normalized to compare across the range of natural variation, infection rate often has the largest effect. Across all plots, bacterial populations which approximately reached their stationary phase density are plotted as 'x's, and bacterial populations that did not reach extinction within 48 hours are plotted as '+'s.

#### Appendix 6. Sensitivity of metrics to inoculum densities

In the main text, we reported how changes in the initial bacterial or phage density (while holding the other constant) can alter peak time on the same order of magnitude as infection rate (Fig 6). Here, we supplement those data. First, we show how changes in initial density while holding the ratio of bacteria to phages constant can alter metrics of bacterial population dynamics (Fig S13). Then, we show that changes in initial density also alter other metrics, like peak density, extinction time, and area under the curve (Figs S14, S15).

Given these results, normalization of initial inoculum densities to be standardized across experiments is essential to infer phage infectivity. However, in practice, it is experimentally impossible to perfectly normalize the initial densities of phages and bacteria. How worried should we be that stochasticity in initial densities will obscure differences in phage infectivity? Here, we addressed the question by quantifying the impact of the two primary sources of noise: uncertainty in estimating bacterial or phage density, and noise from sampling of bacterial or phage suspensions.

First, the density of bacterial cells or phage particles in a suspension is typically estimated by diluting the suspension, plating it, and counting the number of colony or plaque forming units, where this count is a stochastic sample of the true unknown density. Counting protocols can vary in the volume of liquid plated (and therefore the number of colonies or plaques that will be counted) and the number of replicate plates. Here, we used our simulations of growth curves with varying initial densities and infection rate to quantify how much this noise would change the time of peak bacterial density, and then calculate how large of a change in infection rate would produce the same change in time of peak bacterial density. Our results show that noise arising from density estimation will only obscure relatively small differences in infection rate (Fig S16). For instance, a single replicate of 100 colonies limits the noise to obscure differences in infection rate smaller than ±16%, and a single replicate of 100 plaques limits the noise to obscure differences in infection rate smaller than ±11%.

Second, once the density of a suspension has been estimated, only a random sample of that suspension will be inoculated into the well; the actual number of bacteria or phages in that sample is subject to some stochasticity. Here, we used our simulations of growth curves with varying initial densities and infection rate to quantify how much this noise would change the time of peak bacterial density, and then calculate how large of a change in infection rate would produce the same change in time of peak bacterial density. We find that this step produces inconsequential levels of noise, since typical inoculation densities are on the order of $10^4$ – $10^6$ cfu or pfu/mL, and so extremely unlikely to obscure any differences in infection rate (Fig S17).

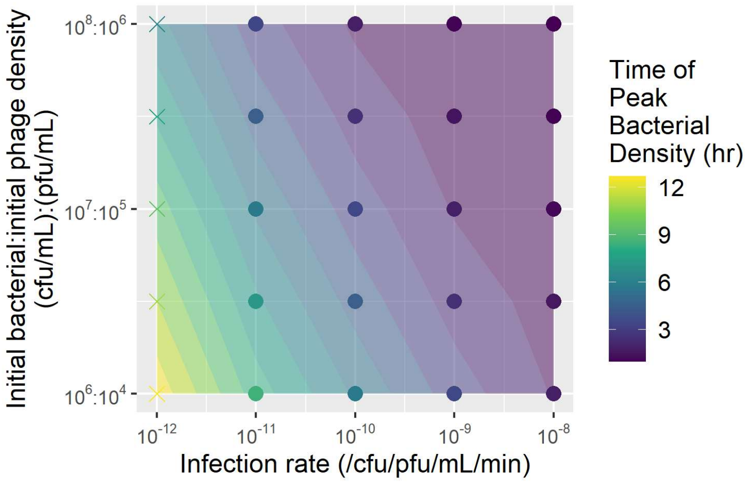


**Figure S13. Inoculum densities with constant MOI alter growth curve metrics.** Bacterial population dynamics were simulated with varying infection rates and co-varying initial bacterial and phage densities, holding the ratio of phages to bacteria (the multiplicity of infection, MOI) constant at 0.01. Each point is a single simulation, with the time when the bacterial density peaked plotted as the color, and linear interpolation of the surface between points for visualization. Growth curves where bacteria approximately reached their stationary phase density are plotted as an "X" here.

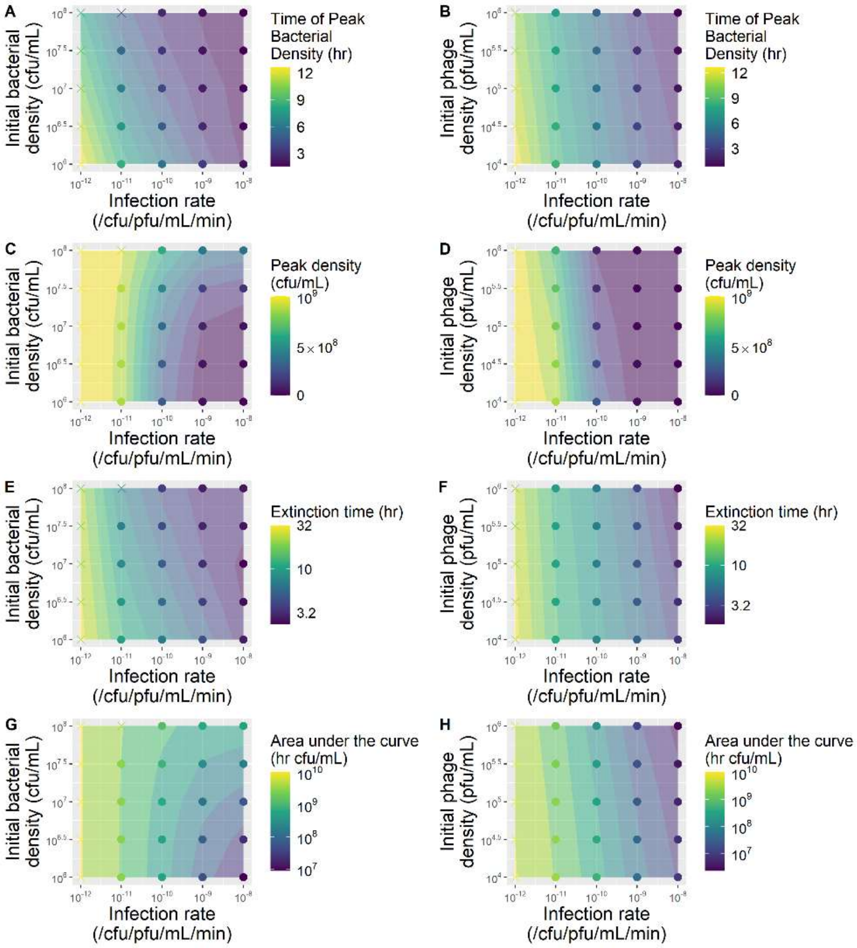


**Figure S14. Inoculum densities alter growth curve metrics.** Bacterial population dynamics were simulated with varying infection rates. **A,C,E,G.** The initial density of phages was held constant at $10^4$ PFU/mL. **B,D,F,H.** The initial density of bacteria was held constant at $10^6$ CFU/mL. Each point is a single simulation, with the time of peak bacterial density (A, B), peak bacterial density (C, D), time when bacterial density dropped below $10^4$ CFU/mL (E, F), or area under the curve (G, H) plotted as the color, and linear interpolation of the surface between points for visualization. Populations where bacteria approximately reached their stationary phase density are plotted as an "X" here.

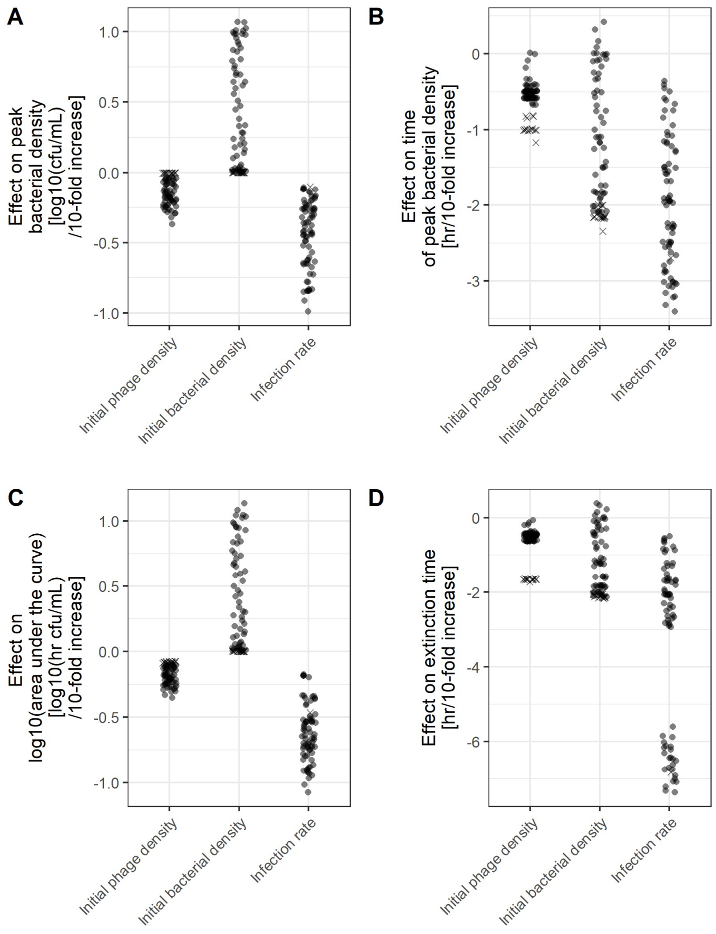


**Figure S15. Metrics of bacterial population dynamics can be sensitive to inoculation densities.** Bacterial population dynamics were simulated with phages with varying infection rates ($10^{-12}$, $10^{-11}$, $10^{-10}$, $10^{-9}$, $10^{-8}$ /CFU/PFU/min), and initial bacterial ($10^{6}$, $10^{6.5}$, $10^{7}$, $10^{7.5}$, $10^{8}$ CFU/mL) and phage ($10^{4}$, $10^{4.5}$, $10^{5}$, $10^{5.5}$, $10^{6}$ PFU/mL) densities. All other parameters were default values (Table 1). We calculated the rate of change of each metric against a 10-fold change in infection rate, initial bacterial density, or initial phage density. **D.** $10^{4}$ CFU/mL was used as the arbitrary threshold to calculate extinction time.

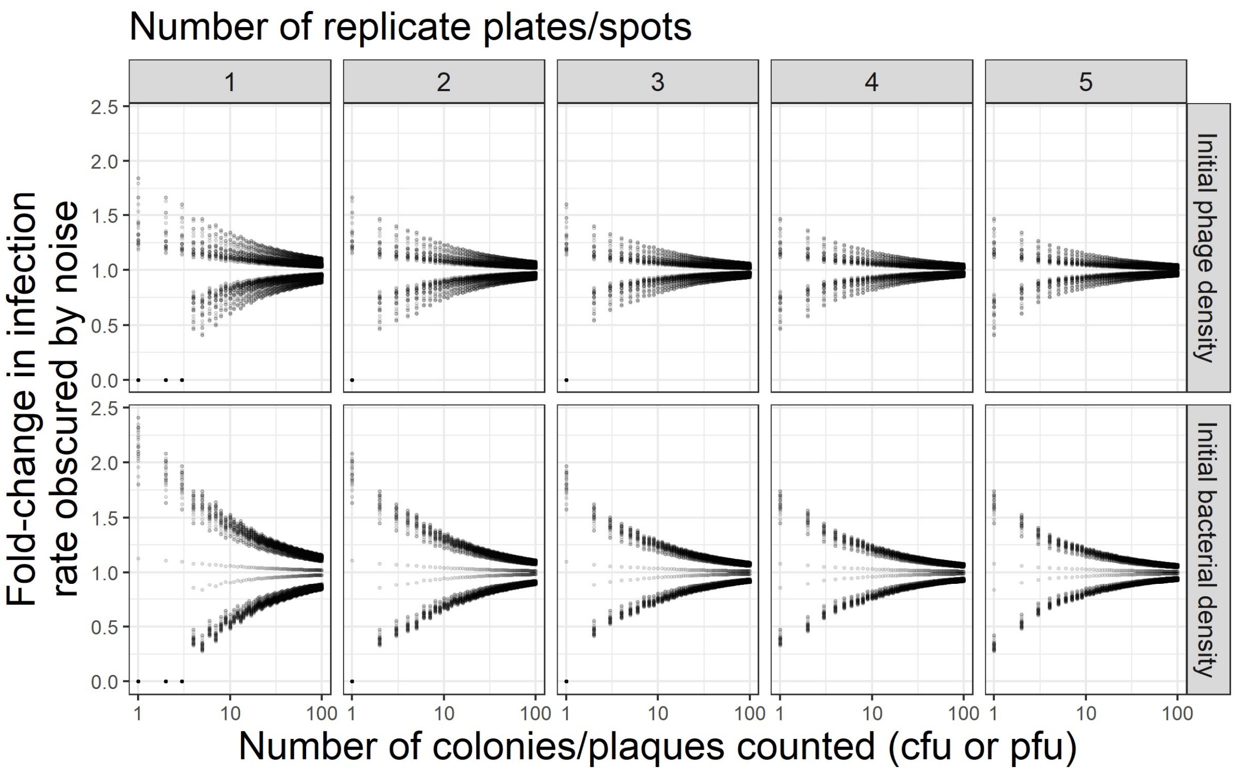


**Figure S16. Density estimation error can obscure small differences in infection rate.** Bacterial population dynamics were simulated with phages with varying infection rates ($10^{-12}$, $10^{-11}$, $10^{-10}$, $10^{-9}$, $10^{-8}$ /CFU/PFU/min), and initial bacterial ($10^{6}$, $10^{6.5}$, $10^{7}$, $10^{7.5}$, $10^{8}$ CFU/mL) and phage ($10^{4}$, $10^{4.5}$, $10^{5}$, $10^{5.5}$, $10^{6}$ PFU/mL) densities. All other parameters were default values (Table 1). We then calculated the rate of change of time of peak bacterial density against changes in infection rate, initial bacterial density, and initial phage density. To compare these values at experimentally realistic scales, we generated estimation noise by calculating the percent error of the 2.5 and 97.5 quantiles of the Poisson distribution with a mean (λ) of number of colonies/plaques × number of replicates. We then calculated how much the generated percent error in initial phage or bacterial density would change the time of peak bacterial density, and then what fold-change in infection rate would be needed produce the same change in time of peak bacterial density.

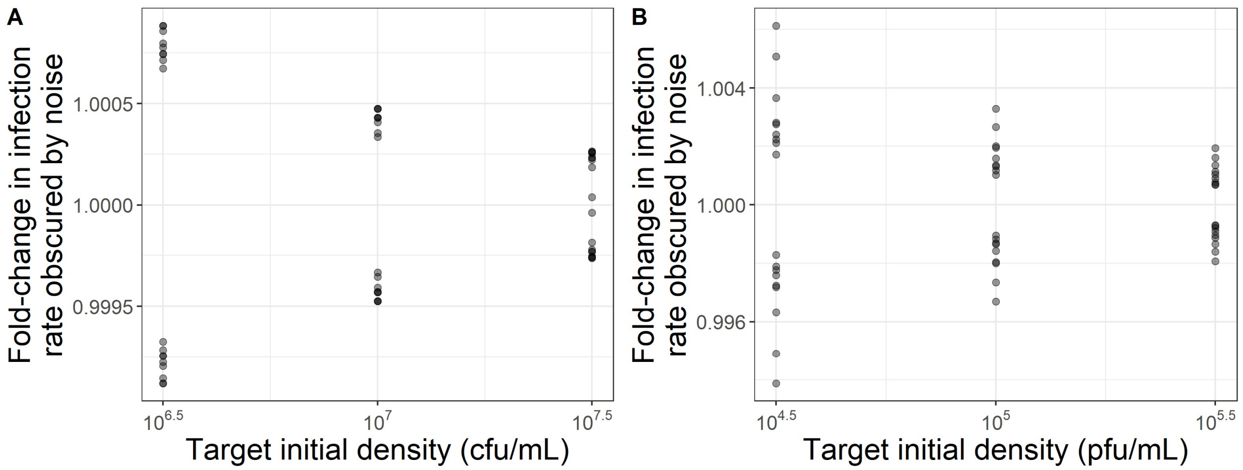


**Figure S17. Sampling error can only obscure inconsequential differences in infection rate.** Bacterial population dynamics were simulated with phages with varying infection rates ($10^{-12}$, $10^{-11}$, $10^{-10}$, $10^{-9}$, $10^{-8}$ /CFU/PFU/min), and initial bacterial ($10^{6}$, $10^{6.5}$, $10^{7}$, $10^{7.5}$, $10^{8}$ CFU/mL) and phage ($10^{4}$, $10^{4.5}$, $10^{5}$, $10^{5.5}$, $10^{6}$ PFU/mL) densities. All other parameters were default values (Table 1). We then calculated the rate of change of time of peak bacterial density against changes in infection rate, initial bacterial density, and initial phage density. To compare these values at experimentally realistic scales, we generated sampling noise by calculating the percent error of the 2.5 and 97.5 quantiles of the Poisson distribution with a mean (λ) of the initial bacterial **(A)** or phage **(B)** density. We then calculated how much the generated percent error in initial phage or bacterial density would change the time of peak bacterial density, and then what fold-change in infection rate would be needed produce the same change in time of peak bacterial density.

#### Appendix 7. Metrics across varying bacterial hosts

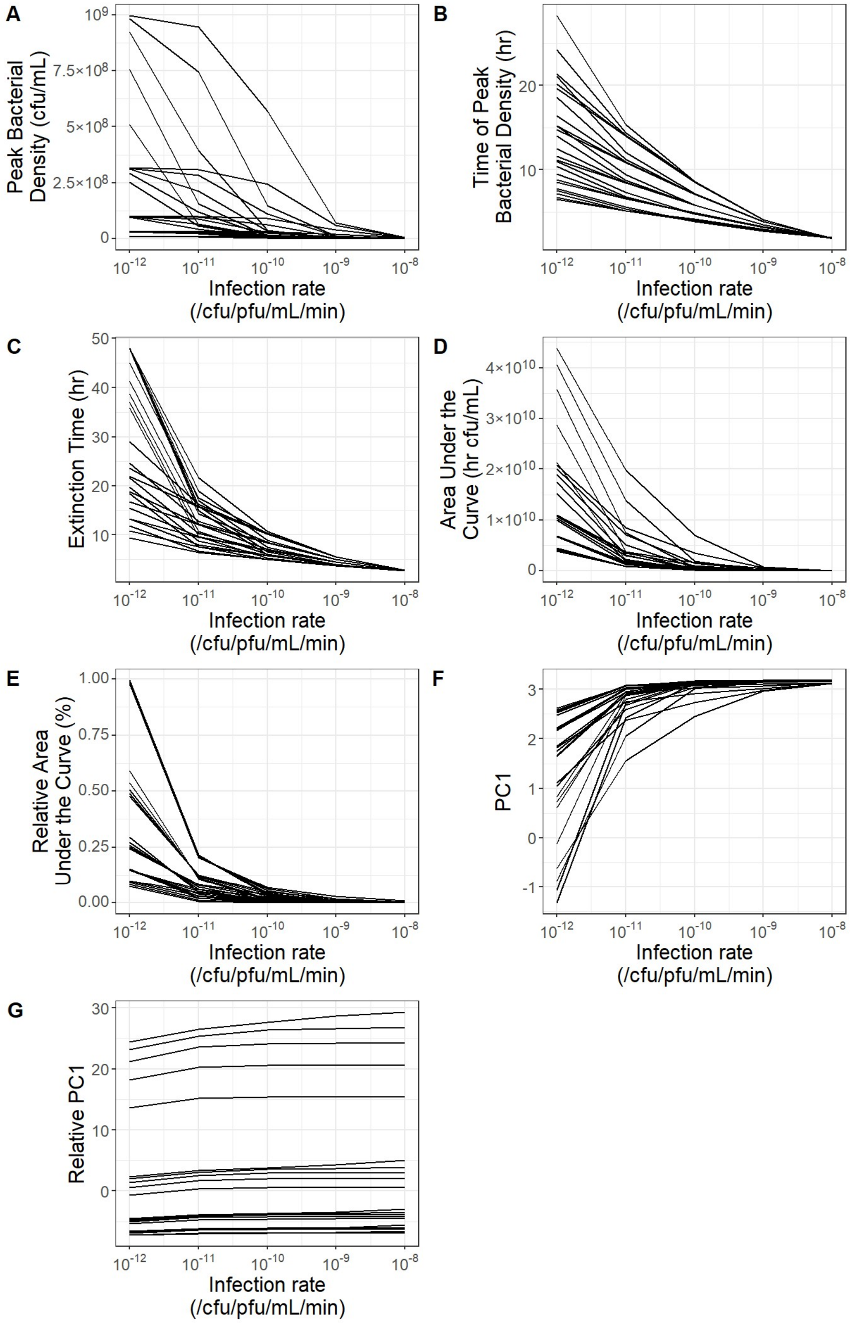


**Figure S18. Metrics of population dynamics generally still correlate with infection rate across bacterial variation.** Bacterial population dynamics were simulated with phages with varying infection rates ($10^{-12}$, $10^{-11}$, $10^{-10}$, $10^{-9}$, $10^{-8}$ /CFU/PFU/min) and bacteria with varying stationary phase densities ($10^{8}$, $10^{8.5}$, $10^{9}$, $10^{9.5}$, $10^{10}$ CFU/mL) and growth rates (0.04, 0.027, 0.018, 0.012, 0.008 /min; doubling times of 17, 26, 39, 58, and 87 mins). Each line plots the metrics calculated from bacterial population dynamics with bacteria having the same growth rate and stationary phase density, across varying infection rates.

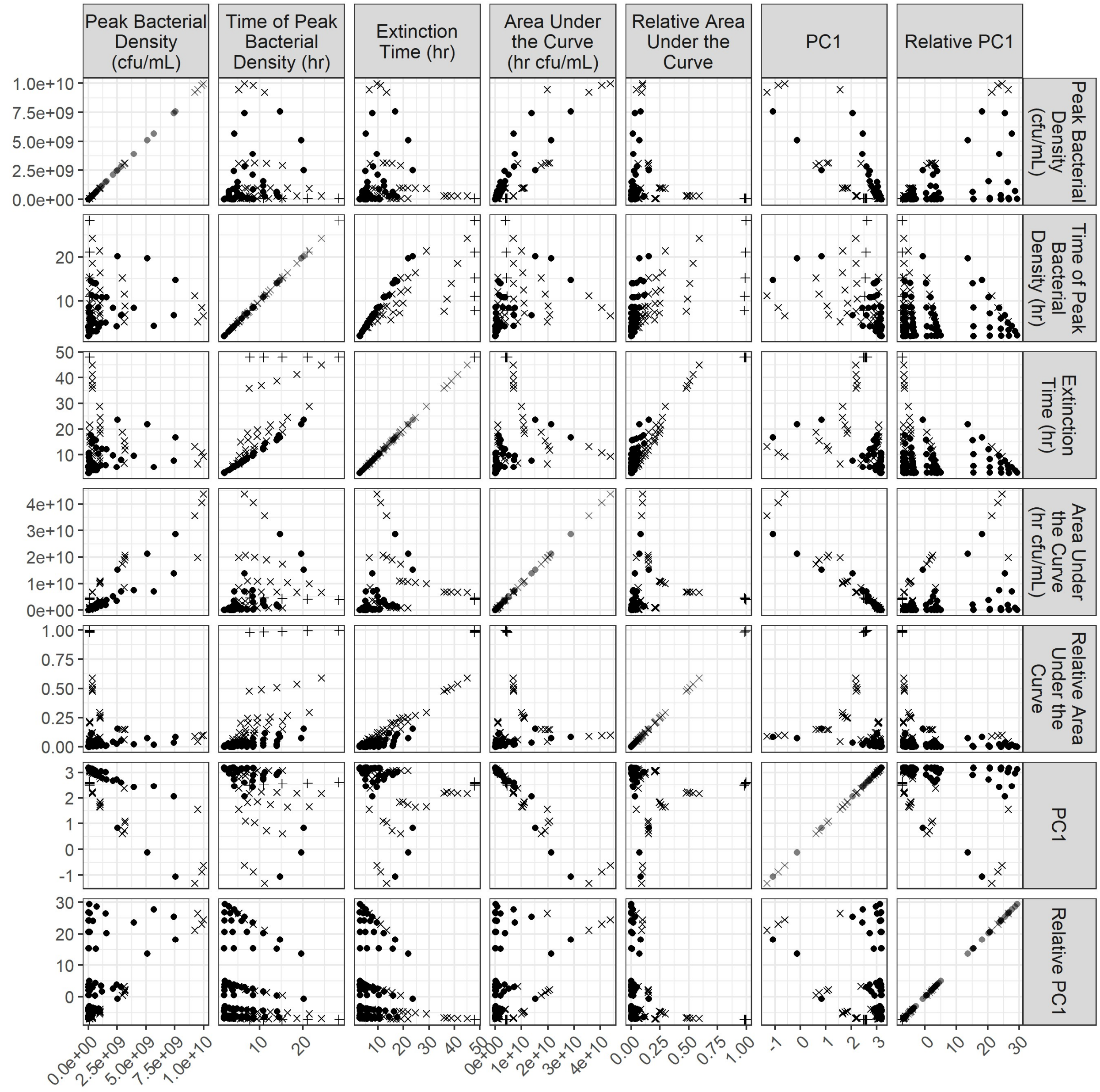


**Figure S19. Correlations between metrics of bacterial population dynamics are weak when bacteria vary in growth rate and stationary phase density.** Bacterial population dynamics were simulated with phages with varying infection rates ($10^{-12}$, $10^{-11}$, $10^{-10}$, $10^{-9}$, $10^{-8}$ /CFU/PFU/min) and bacteria with varying stationary phase densities ($10^{8}$, $10^{8.5}$, $10^{9}$, $10^{9.5}$, $10^{10}$ CFU/mL) and growth rates (0.04, 0.027, 0.018, 0.012, 0.008 /min; doubling times of 17, 26, 39, 58, and 87 mins). Bacterial populations which approximately reached their stationary phase density are plotted as ‘x’s, and bacterial populations that did not reach extinction within 48 hours are plotted as ‘+’s. PC1 is the first principal component from a principal component analysis of the bacterial population dynamics.

Some proposed methods for quantifying infectivity by normalizing area under the curve (AUC) involve comparing relative AUC across varying initial multiplicities of infection (44) (where relative AUC is AUC in the presence of a phage divided by AUC in the absence of a phage). They define a "Virulence Index" which is simply 1 – relative AUC. However, it remains unclear what patterns we should expect from this Virulence Index. Here, we find that Virulence Index increases approximately linearly with log initial MOI across phages which vary in infection rate, largely regardless of bacterial growth rate or stationary phase density (Fig S20).

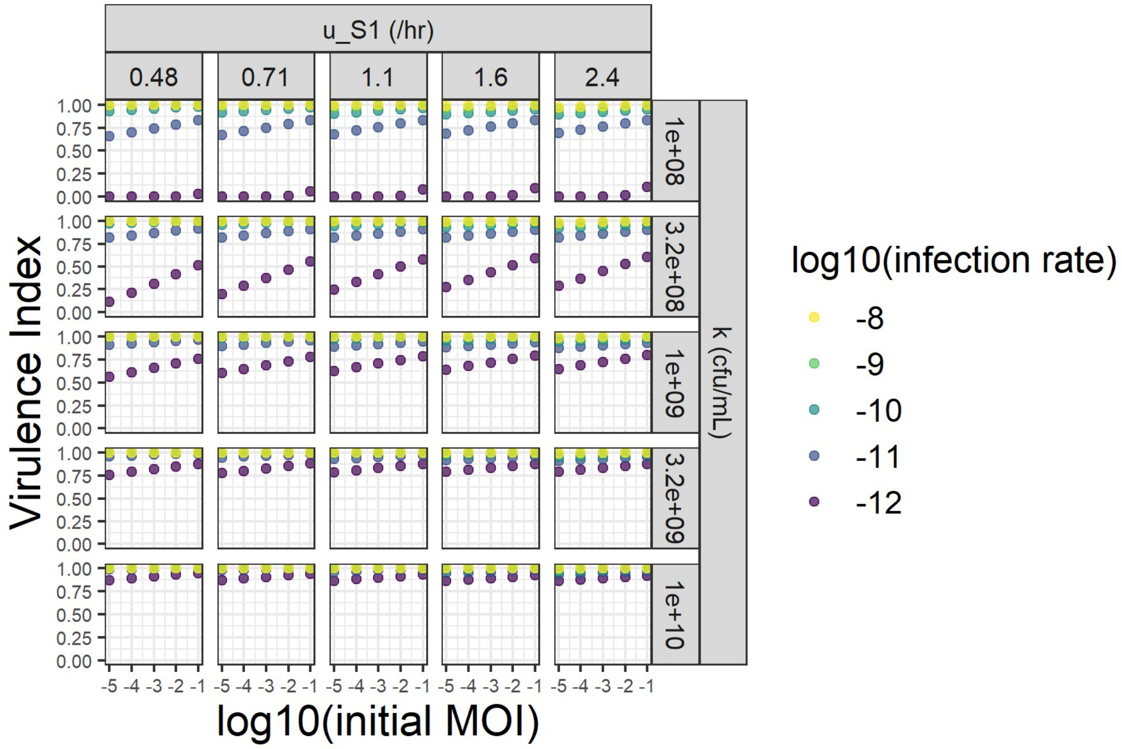


**Figure S20. Relative area under the curve (AUC) is approximately linear with initial multiplicity of infection.** The "Virulence Index" is defined by (44) as 1 – Relative AUC. Shown are the virulence index values for growth curves of phages varying in their infection rate on bacteria which vary in their growth rate (u_S1) and stationary phase density (k). Here, we show that the Virulence Index increases approximately linearly with the initial multiplicity of infection (MOI) of a growth curve assay.

#### Appendix 8. Empirical growth curves

In this paper we have used mathematical models to thoroughly explore how bacterial population dynamics with phages can be used to infer infectivity. This theoretical exploration was originally motivated by experiments we carried out with phage Phi2 and its host *Pseudomonas fluorescens* (87), as well as the extensive body of existing empirical papers assessing how bacterial population dynamics might be used to infer phage infectivity (37–50). For our experiments, we took bacterial isolates that vary in their susceptibility to Phi2 infection because of evolved resistance from a previous study (88). In that study, bacteria were evolved in environments which differed in their spatial distribution of phages (control, local, or global), and in media and temperatures where phages grew better or worse ('strong' or 'weak'). We selected the ancestor and nine evolved bacterial strains that spanned a wide range of susceptibility to infection by Phi2 to use as a set of strains to assess patterns in phage-bacteria growth curves.

To collect bacterial population dynamics, bacteria were grown overnight with shaking at 28°C in 10 mL of King's B media (0 g/L LP0037 Oxoid Bacteriological Peptone, 15 g/L glycerol, 1.5 g/L potassium phosphate, and 0.6 g/L magnesium sulfate). Overnight cultures were then inoculated into 200 µL of media to achieve an initial density of $10^5$ CFU/mL. For each isolate, duplicate wells were inoculated for bacteria-alone growth, and triplicate wells were inoculated with bacteria and wild-type Phi2 at an initial multiplicity of infection of 0.1. Plates were incubated in an automated spectrophotometer at 28°C for 24 hours with regular readings of the optical density at 600 nm. This entire experiment was repeated for a total of three batches. Data was analyzed using gcplyr (63) in R v4.2.2. All data and code used to analyze and visualize data is available at https://github.com/mikeblazanin/growth-curves.

We observe that our theoretical results reproduce a notable pattern from our empirical results: the exponential phase of bacterial growth provides little information about phage infectivity (Fig S21, reproducing patterns from Fig 1). In contrast, later phases of bacterial population dynamics reflect susceptibility to phage infection. For instance, metrics like peak density, peak time, and area under the curve all correlate well with a traditional measure of susceptibility to phage (efficiency of plaquing, EOP, Fig S22). Interestingly, EOP is itself a complex measure with poorly-understood relationships to phage traits like burst size, lysis time, and infection rate (72), an avenue for further theoretical and empirical work.

However, we also noticed deviations from our theoretical outcomes. For instance, the maximum rate of population decline was correlated with phage infectivity in our experimental data (Fig S22), in contrast to our theoretical findings (Fig 2). Additionally, and most notably, many of the moderately-susceptible bacteria had growth curves with a pattern of partial population lysis, where the bacterial population rose, peaked, and then declined partially (Fig S21). These observations motivated our exploration of additional model structures (Fig 8, Appendix 9).

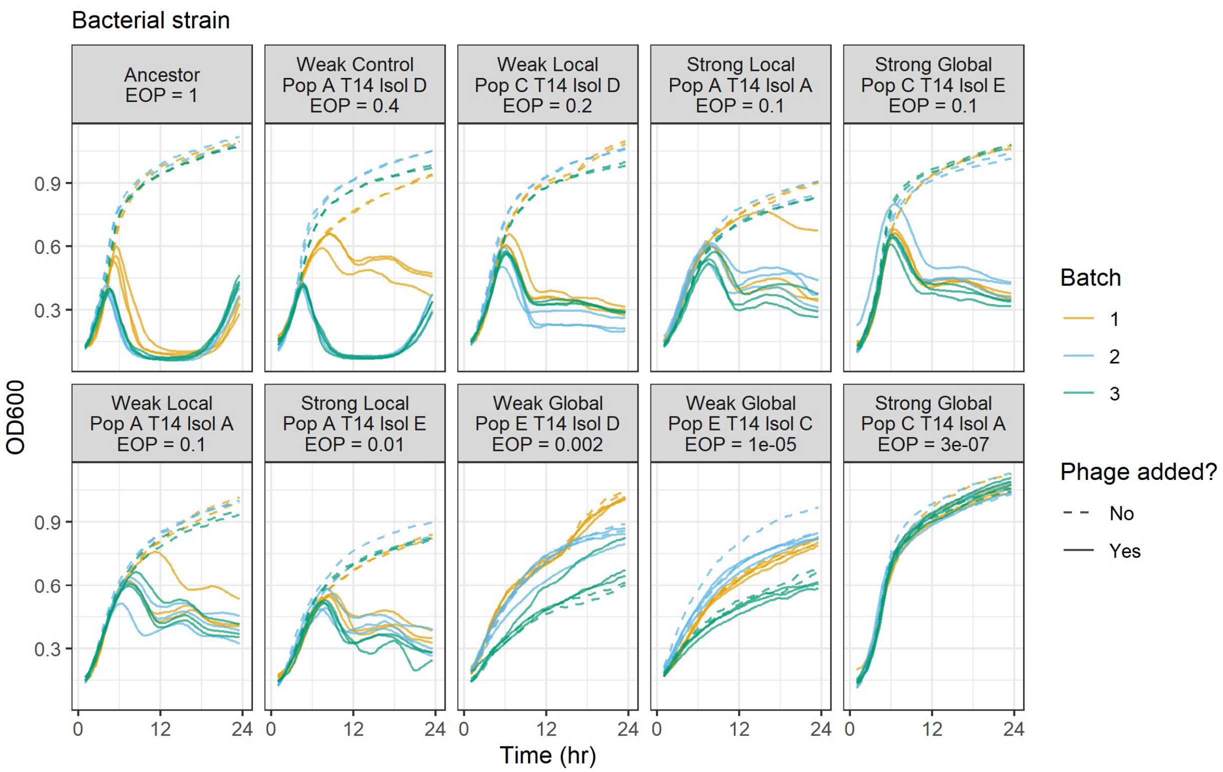


**Figure S21. Experimental phage-bacteria growth curves exhibit similarity and differences to theoretical growth curves.** We conducted growth curves with phage Phi2 and *Pseudomonas fluorescens*, using the ancestral and 9 evolved bacterial mutants from (88). We collected three technical replicates with phages added and two technical replicates with bacteria alone across three difference batches. Efficiency of plaquing (EOP) data from (88) is a standard measure of susceptibility to phage infection relative to a reference strain (here the Ancestor). These experimental curves qualitatively reproduce a pattern found in simulated growth curves: that the initial growth period of the bacteria provides little information on phage infectivity. They also prompted the development of additional models that could explain the observed partial lysis of bacterial populations seen with many of the bacterial isolates.

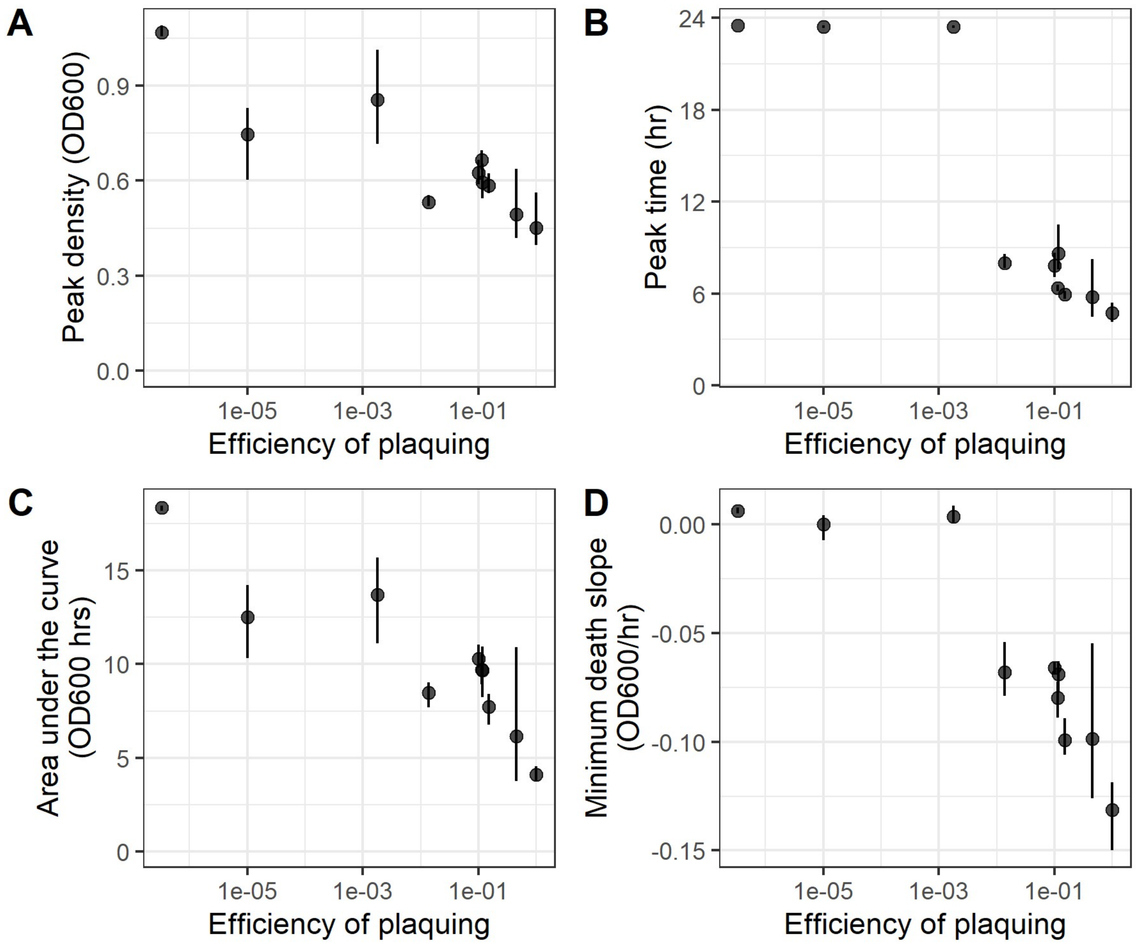


**Figure S22. Metrics of experimental phage-bacteria growth curves correlate well with traditional measures of susceptibility to phages.** We conducted growth curves with phage Phi2 and *Pseudomonas fluorescens*, using the ancestral and 9 evolved bacterial mutants from (88). Shown are the average of three batches, with lines denoting the range of the three batches. Efficiency of plaquing (EOP) data from (88) is a standard measure of susceptibility to phage infection relative to a reference strain (here the Ancestor).

#### Appendix 9. Metrics across the longer term with plasticity or evolution

Here, we expand our exploration of how bacterial population dynamics can be used to infer the effects of phages over timescales where bacterial resistance can change, due to plasticity or evolution. Specifically, we use previously-published approaches (51) to model three scenarios: phage growth weakening as bacterial growth slows (20, 51, 57, 60, 64, 65), bacterial cells transitioning into a phenotypically resistant state (61), and bacteria evolving mutations that confer resistance to phage infection.

##### Continuous approximation model for all three scenarios

Thus far, we have used a model where phages lyse their infected hosts a precise, discrete amount of time after they infected the cell. While this model produces simulations which can quite precisely recreate the life cycle of phages, it can be difficult to evaluate analytically. In order to facilitate analytical evaluation of our model, we derived a simple ordinary differential equation version of the delay differential model we have used so far [Eqs. S1 – S4]. Specifically, rather than cells lysing a finite delay after they became infected, there are $n_I$ stages of infection that cells proceed through at rate 1/τ before lysing and releasing new phage particles. Past work has shown that, as $n_I$ increases, this model increasingly resembles the original delay differential system (20, 51, 65).

$$\frac{dS}{dt} = u_S S \frac{N}{k} - aSP \tag{S1}$$

$$\frac{dI_1}{dt} = aSP - \frac{n_I I_1}{\tau} \tag{S2}$$

$$\frac{dI_j}{dt} = \frac{n_I I_{j-1}}{\tau} - \frac{n_I I_j}{\tau} \; for\; j \in 2 \ldots n_I \tag{S3}$$

$$\frac{dP}{dt} = -aSP + b\frac{n_I I_{n_I}}{\tau} \tag{S4}$$

We then used versions of this model to determine how each scenario altered the possible outcomes of bacterial population dynamics.

##### Plasticity in susceptibility to phage

In the first scenario, we model plastic changes in the infection rate (60), burst size (57, 60, 64), and lysis time (20, 51, 65) as nutrients become depleted.

For *simulations* of changes in infection rate and burst size, we can implement these changes in the original delay differential model:

$$\frac{dS}{dt} = uS\frac{N}{k} - a_t SP \tag{S5}$$

$$\frac{dI}{dt} = a_t SP - a_{t-\tau} S_{t-\tau} P_{t-\tau} \tag{S6}$$

$$\frac{dP}{dt} = -a_t SP + b_t a_{t-\tau} S_{t-\tau} P_{t-\tau} - a_t zIP \tag{S7}$$

$$\frac{dN}{dt} = -uS\frac{N}{k} + da_{t-\tau}S_{t-\tau}P_{t-\tau} \quad [S8]$$

Where:

$$a_t = a * \max\left(0, 1 - f_a + f_a\frac{N_t}{k}\right) \quad [S9]$$

$$b_t = b * \max\left(0, 1 - f_b + f_b\frac{N_t}{k}\right) \quad [S10]$$

Here, $a_t$ and $b_t$ decline linearly with decreasing nutrient availability. The slope of that decline, and the total decrease when nutrients have been completely depleted, is controlled by the *f* parameter.

However, the original delay differential model only works for *simulations* of changes in infection rate and burst size. For analytical evaluation of changes in infection rate and burst size, and both analytical evaluation and simulation of changes in lysis time, we use the ordinary differential equation version of the model [Eqs. S1 – S4] with some modifications:

$$\frac{dS}{dt} = u_S S\frac{N}{k} - a_t SP \quad [S11]$$

$$\frac{dI_1}{dt} = a_t SP - \frac{n_I I_1}{\tau_t} \quad [S12]$$

$$\frac{dI_j}{dt} = a_t SP - \frac{n_I I_j}{\tau_t} \; for\; j \in 2 \ldots n_I \quad [S13]$$

$$\frac{dP}{dt} = -a_t SP + b_t\frac{n_I I_{n_I}}{\tau_t} \quad [S14]$$

Where:

$$a_t = a * \max\left(0, 1 - f_a + f_a\frac{N_t}{k}\right) \quad [S15]$$

$$b_t = b * \max\left(0, 1 - f_b + f_b\frac{N_t}{k}\right) \quad [S16]$$

$$\tau_t = \frac{\tau}{\max\left(0, 1 - f_\tau + f_\tau\frac{N_t}{k}\right)} \quad [S17]$$

#### **Transitions into resistant state (by plasticity or mutation)**

The second and third scenarios involve bacterial cells transitioning into a resistant sub-population, either via some phenotypic plasticity or an evolutionary mutation (61). Here, we focus on a simple model with two subpopulations: one which is susceptible to phage infection, and one which is completely resistant to phage infection. Moreover, we only model transitions from the first subpopulation to the second, at a rate h relative to the bacterial growth rate.

For the second scenario, these transitions were modeled in three ways: constant, increasing as nutrients become depleted, and decreasing as nutrients become depleted. For the third scenario, we only modeled these transitions as being constant, since per-generation mutation rate is not expected to change strongly based on the current growth rate.

<u>Constant transitions to resistance</u>

$$\frac{dS}{dt} = u_S S \frac{N}{k} - a_t SP - hu_S S \quad [S18]$$

$$\frac{dR}{dt} = u_R R \frac{N}{k} + hu_S S \quad [S19]$$

$$\frac{dN}{dt} = -u_S S \frac{N}{k} - u_R R \frac{N}{k} \quad [S20]$$

<u>Increasing transitions to resistance as nutrients become depleted</u>

$$\frac{dS}{dt} = u_S S \frac{N}{k} - a_t SP - hu_S S \left(1 - \frac{N}{k}\right) \quad [S21]$$

$$\frac{dR}{dt} = u_R R \frac{N}{k} + hu_S S \left(1 - \frac{N}{k}\right) \quad [S22]$$

$$\frac{dN}{dt} = -u_S S \frac{N}{k} - u_R R \frac{N}{k} \quad [S23]$$

<u>Decreasing transitions to resistance as nutrients become depleted</u>

$$\frac{dS}{dt} = u_S S \frac{N}{k} - a_t SP - hu_S S \frac{N}{k} \quad [S24]$$

$$\frac{dR}{dt} = u_R R \frac{N}{k} + hu_S S \frac{N}{k} \quad [S25]$$

$$\frac{dN}{dt} = -u_S S \frac{N}{k} - u_R R \frac{N}{k} \quad [S26]$$

#### <u>Analytical evaluation</u>

We then used these models (Equations S5 – S10, S11 – S17, S18 – S20, S21 – S23, S24 – S26) to evaluate possible outcomes of bacterial population dynamics and identify all possible equilibria (Table S2). All models had an equilibrium when susceptible and infected bacteria had gone extinct, with models where bacteria could transition into a resistant state only reaching this equilibrium if resistant cells could not grow or once nutrients were entirely depleted. Many models also had an equilibrium where phages were absent or had gone extinct. Notably, models with plasticity in infection rate, burst size, or lysis time have a separatrix at $N = \frac{1}{f}$ that separates whether phages or susceptible bacteria will go extinct. If the bacterial population can consume the available nutrients below $\frac{1}{f}$ before being killed off by the phages, the phage growth rate will become zero and the bacterial population will persist; otherwise, if the bacterial population cannot consume the nutrients below $\frac{1}{f}$ fast enough, the phage population will drive them extinct.

**Table S2. Equilibria of models with plasticity or evolution.** Each version of the model is shown in the first column, with equilibria conditions listed as the headers of additional columns. When the equilibria hold for the model with no additional conditions an 'x' is used, otherwise the additional necessary and sufficient conditions are listed in the table.

| | *Equilibria* | | | *Special case equilibria* |
|---|---|---|---|---|
| **Case** | **S = 0**<br>**I = 0** | **I = 0**<br>**N = 0** | **S = 0**<br>$\frac{1}{f}+\frac{N}{k}\leq 1$ | **P = 0**<br>**N = 0**<br>**f ≥ 1** |
| **Base model**<br>**[No plasticity or evolution]** | x | P = 0 | | |
| **Plastic $a_t$** | x | P = 0<br>or<br>f ≥ 1 | | |
| **Plastic $b_t$** | x | P = 0 | | |
| **Plastic $\tau_t$** | x | P = 0 | x | x |
| $u_R = 0$ **and constant transition rate to resistance** | x | | | |
| $u_R = 0$ **and decreasing transition rate to resistance with nutrient scarcity** | x | P = 0 | | |
| $u_R = 0$ **and increasing transition rate to resistance with nutrient scarcity** | x | | | |
| $u_R > 0$ **and constant transition rate to resistance** | R = 0<br>or<br>N = 0 | | | |
| $u_R > 0$ **and decreasing transition rate to resistance** | R = 0<br>or<br>N = 0 | R = 0 | | |
| $u_R > 0$ **and increasing transition rate to resistance** | R = 0<br>or<br>N = 0 | | | |

#### Simulations of plasticity in susceptibility to phage

In the main text, we show a subset of simulations with plasticity in infection rate (Fig 8A, 8D). Here, we show a broader range of parameter values for plasticity in infection rate (Fig S23), as well as simulations with plasticity in burst size (Fig S24) and lysis time (Fig S25). Table S3 lists definitions and values for parameters in these simulations that cannot be found in the main text (Table 1).

**Table S3.** Parameters and definitions for terms exclusive to supplemental equations. All other parameters used values listed in Table 1. Unless otherwise noted, all simulations used the default value listed. For the noted exceptions, simulations typically used the range of values in brackets.

| Parameter | Definition | Default Value [Range of values] |
|---|---|---|
| $f_b$ | Degree of burst size plasticity | 0 [0 – 3] |
| $f_\tau$ | Degree of lysis rate plasticity | 0 [0 – 3] |
| $n_I$ | Number of infected cell stages | 50 |

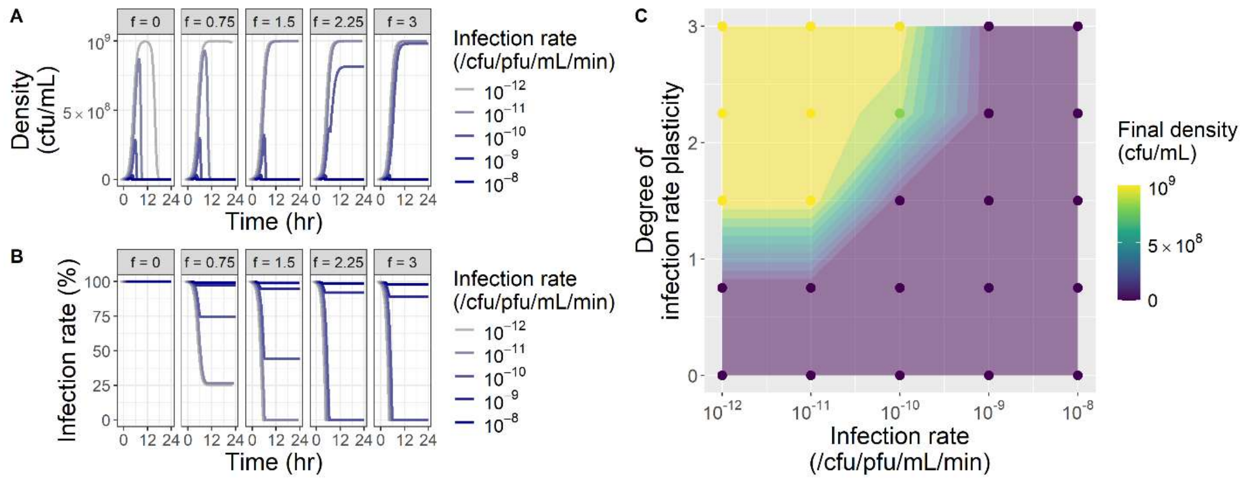


**Figure S23. Plasticity in infection rate can alter the outcome of bacterial population dynamics.** Bacterial population dynamics were simulated with phages with varying infection rates and varying degrees of plasticity in infection rate, where infection rate declines linearly with declining bacterial growth rate. Bacterial populations that deplete nutrients enough that the phage infection rate reaches 0 (shown in B) can persist without ever crashing. Curves in A and B have been slightly offset horizontally for visualization.

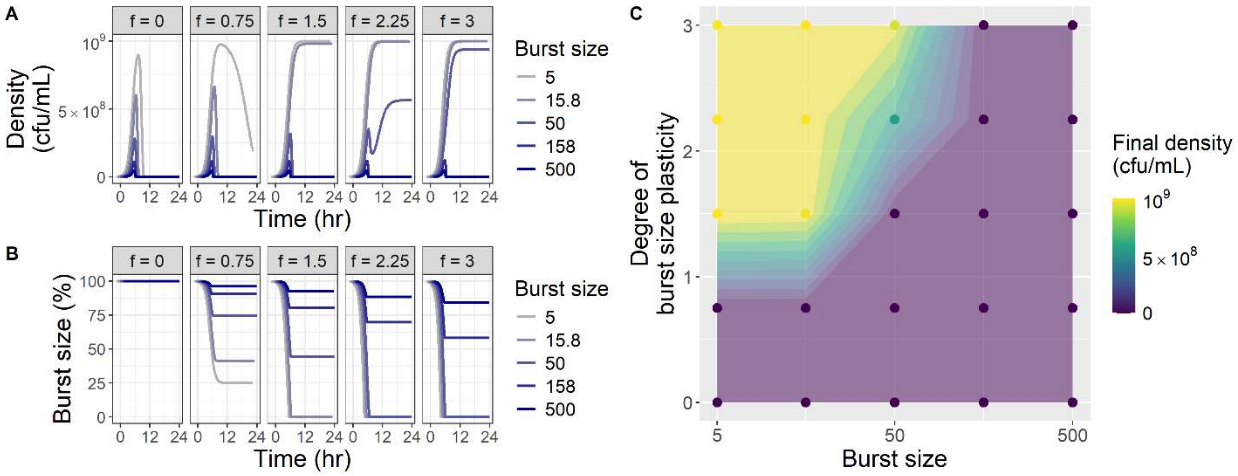


**Figure S24. Plasticity in burst size can alter the outcome of bacterial population dynamics.** Bacterial population dynamics were simulated with phages with varying burst sizes and varying degrees of plasticity in burst size, where burst size declines linearly with declining bacterial growth rate. Bacterial populations that deplete nutrients enough that the phage burst size reaches 0 (shown in B) can persist without ever crashing. Curves in A and B have been slightly offset horizontally for visualization.

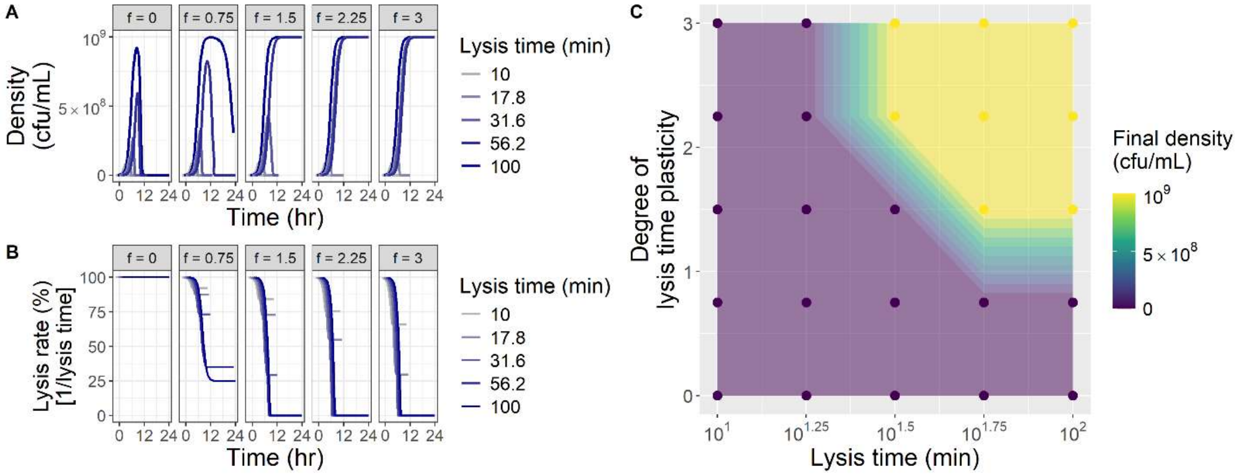


**Figure S25. Plasticity in lysis time can alter the outcome of bacterial population dynamics.** Bacterial population dynamics were simulated with phages with varying lysis rates (the inverse of lysis time) and varying degrees of plasticity in lysis rate, where lysis rate declines linearly with declining bacterial growth rate (Eqs. S11-S14, S17, with $n_l$= 50). Bacterial populations that deplete nutrients enough that the phage burst size reaches 0 (shown in B) can persist without ever crashing. Curves in A and B have been slightly offset horizontally for visualization.

#### Simulations of plastic transitions into resistant state

In the main text, we show a subset of simulations with transitions into a phenotypically-resistant state (Fig 8B, 8E). Here, we show a broader range of parameter values (Fig S26). We also show how debris from lysed cells can cause similar patterns of partial population lysis when density is measured optically (Fig S27).

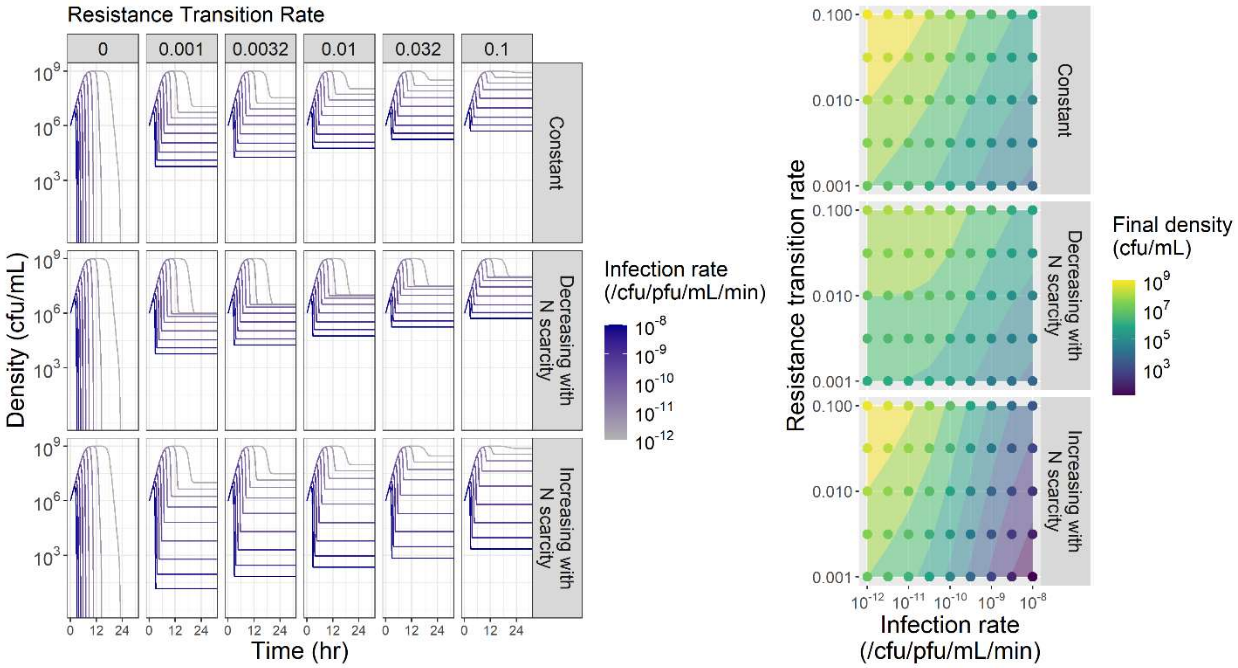


**Figure S26. Transitions into resistant state can produce patterns of partial population lysis.** Bacterial population dynamics were simulated with phages with varying infection rates, where bacteria transitioned into a non-growing resistant state at varying rates that could depend on nutrient availability (Eqs. S18 – S26).

Patterns of partial population lysis can also be produced by debris when density is measured optically (e.g. by absorbance). To test the potential role of debris, we simply define an additional population of debris that grows every time an infected cell lyses (Eq. S27), with a conversion factor $c$ that determines how much debris is generated from each cell lysis.

$$\frac{dD}{dt} = cS_{t-\tau}P_{t-\tau} \qquad \text{[S27]}$$

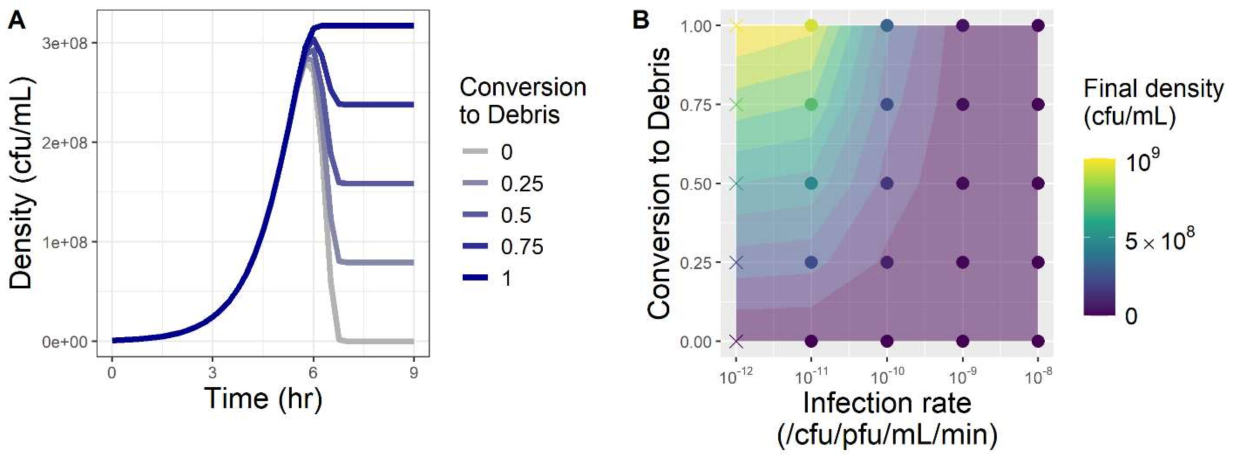


**Figure S27. Debris from lysed calls can produce patterns of partial population lysis.** Bacterial population dynamics were simulated with phages with varying infection rates, where bacteria lysed by phages were converted to debris which continued to contribute to density measures at varying rates.

#### <u>Simulations of evolution into resistant state</u>

In the main text, we show a subset of simulations with mutations conferring resistance (Fig 8C, 8F) where no nutrients are returned to the environment on cell lysis, and where all resistant mutants arise during the course of the simulation. Here, we show how the return of nutrients to the environment by cell lysis as well as the initial frequency of resistant mutants can alter the re-emergence time of the bacterial population (Fig S28). We also show that peak density, the time of peak density, and (when calculable) extinction time remain good indicators of phage infectivity alone, independent of the effects of plasticity or evolution (Fig S29).

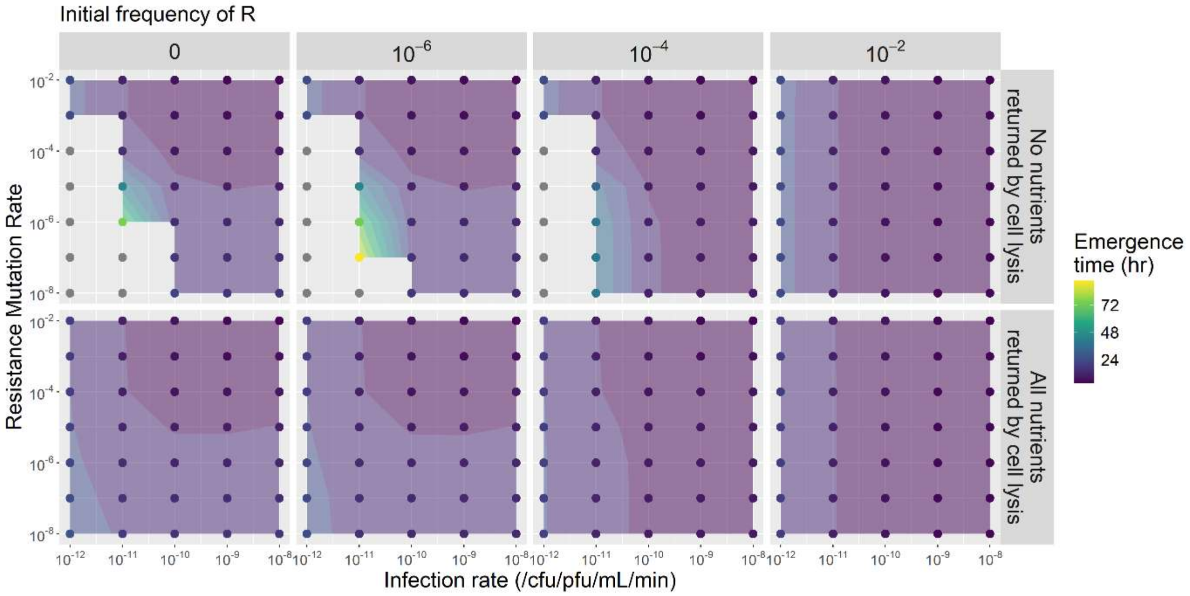


**Figure S28. Evolution can alter the shape of bacterial population dynamics over longer timescales.** Bacterial population dynamics were simulated with phages with varying infection rates, where bacteria evolved cost-free mutations providing complete phage resistance at varying rates, nutrients were or were not returned to the environment by cell lysis, and the frequency of resistant bacteria in the initial population varied.

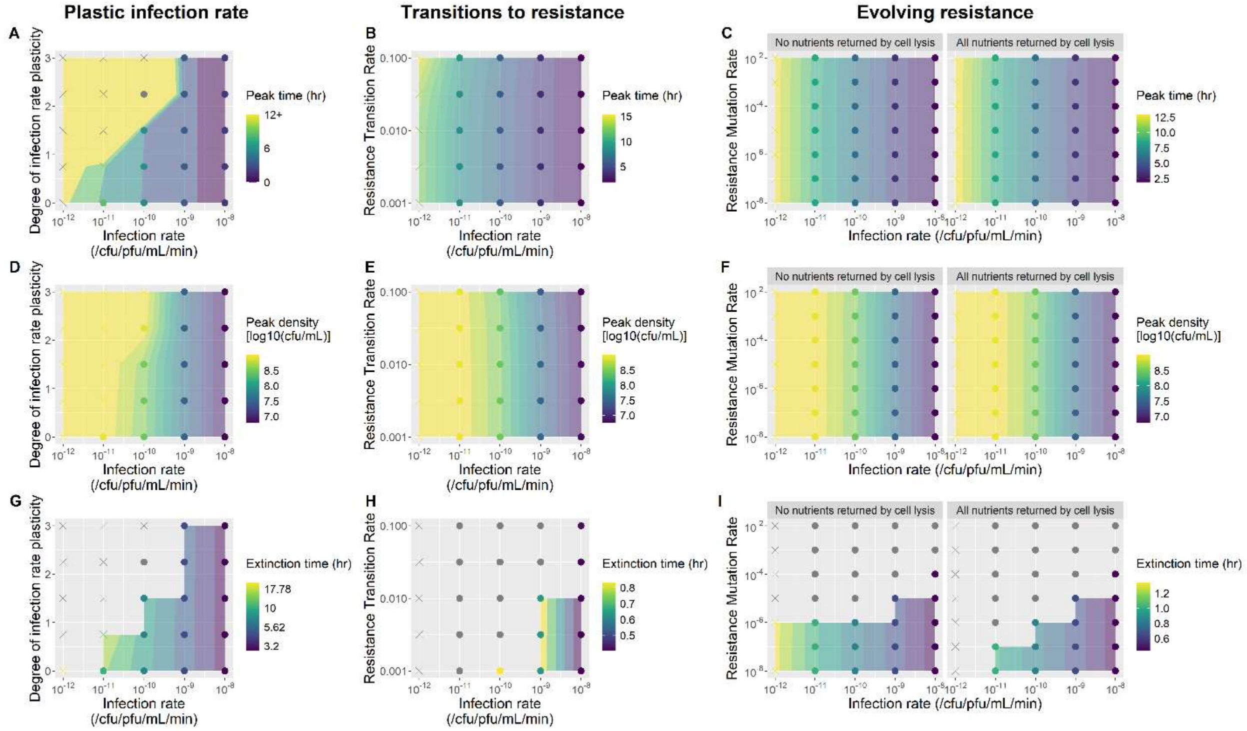

**Figure S29. Metrics of peak density, time of peak density, and extinction time are good indicators of phage infectivity alone, even when bacteria have plasticity or evolution.** Bacterial population dynamics were simulated with phages with varying infection rates ($10^{-12}$, $10^{-11}$, $10^{-10}$, $10^{-9}$, $10^{-8}$ /CFU/PFU/min). **A,D,G.** Phages were simulated with varying degrees of plasticity in infection rate, where infection rate declines linearly with declining bacterial growth rate. **B,E,H.** Bacteria were simulated with transitions into a non-growing resistant state at varying rates that were constant across time. **C,F,I.** Bacteria were simulated with the evolution of cost-free mutations providing complete phage resistance at varying rates, and nutrients were or were not returned to the environment by cell lysis. **H.** $10^{5.99}$ was used as the arbitrary threshold for extinction time here ($10^4$ was used in all other panels).
